# Repetitive head impacts induce neuronal loss and neuroinflammation in young athletes

**DOI:** 10.1101/2024.03.26.586815

**Authors:** Morgane L.M.D. Butler, Nida Pervaiz, Kerry Breen, Samantha Calderazzo, Petra Ypsilantis, Yichen Wang, Julia Cammasola Breda, Sarah Mazzilli, Raymond Nicks, Elizabeth Spurlock, Marco M. Hefti, Kimberly L. Fiock, Bertrand R. Huber, Victor E. Alvarez, Thor D. Stein, Joshua D. Campbell, Ann C. McKee, Jonathan D. Cherry

## Abstract

Repetitive head impacts (RHI) sustained from contact sports are the largest risk factor for chronic traumatic encephalopathy (CTE). Currently, CTE can only be diagnosed after death and the multicellular cascade of events that trigger initial hyperphosphorylated tau (p-tau) deposition remain unclear. Further, the symptoms endorsed by young individuals with early disease are not fully explained by the extent of p-tau deposition, severely hampering development of therapeutic interventions. Here, we show that RHI exposure associates with a multicellular response in young individuals (<51 years old) prior to the onset of CTE p-tau pathology that correlates with number of years of RHI exposure. Leveraging single nucleus RNA sequencing of tissue from 8 control, 9 RHI-exposed, and 11 low stage CTE individuals, we identify SPP1+ inflammatory microglia, angiogenic and inflamed endothelial cell profiles, reactive astrocytes, and altered synaptic gene expression in excitatory and inhibitory neurons in all individuals with exposure to RHI. Surprisingly, we also observe a significant loss of cortical sulcus layer 2/3 neurons in contact sport athletes compared to controls independent of p-tau pathology. Finally, we identify TGFB1 as a potential signal mediating microglia-endothelial cell cross talk through ligand-receptor analysis. These results provide robust evidence that multiple years of RHI exposure is sufficient to induce lasting cellular alterations that may underlie p-tau deposition and help explain the early pathogenesis in young former contact sport athletes. Furthermore, these data identify specific cellular responses to repetitive head impacts that may direct future identification of diagnostic and therapeutic strategies for CTE.

## Introduction

Each year, millions of individuals are exposed to repetitive head impacts (RHI) through contact sports, military service, and domestic violence. These RHIs are often non-symptomatic, non-concussive, and can occur thousands of times per year, over the course of decades in some cases. Chronic traumatic encephalopathy (CTE), a progressive tauopathy caused by exposure to RHI, is observed in individuals as young as 17^1,2^. Risk for CTE in exposed individuals is associated with the number of years of exposure to RHI and the cumulative force of the hits endured^3,4^. While much of the current research is focused on severe CTE in older individuals, a recent case series of 152 brains from donors under the age of 30 identified 63 brains with CTE, highlighting that RHI-driven disease is also pressing concern in the young population^2^. Currently, CTE can only be diagnosed postmortem through identification of hyperphosphorylated tau (p-tau) aggregates in neurons around blood vessels at the depth of the cortical sulcus. Our previous research suggests that microglia-mediated neuroinflammation occurs prior to the deposition of p-tau^5^. Additionally, other work has demonstrated RHI exposure is associated with astrocytic activation, white matter inflammation and damage, blood-brain barrier (BBB) breakdown, serum protein leakage, and increases in vascular density in the CTE brain^5–9^. These cellular changes occur prior to overt neurodegeneration and are likely driving many of the early clinical impairments not explained by the occurrence and extent of p-tau pathology. However, studies examining the full extent of these cellular phenotypes have been limited. A detailed characterization of the early cellular changes in young RHI-exposed athletes is necessary to understand the pathogenic mechanisms in CTE and to identify novel biomarkers or therapeutic targets relevant to early disease stages.

## Results

### Cell type identification and cell proportion analysis across pathological groups

To identify the earliest RHI driven changes, we performed single nucleus RNA sequencing (snRNAseq) using autopsy-confirmed frozen human brain tissue from 28 young individuals. 8 non RHI-exposed controls, 9 RHI-exposed individuals without CTE pathology, and 11 RHI-exposed individuals with diagnosed CTE stage 1 or 2 were included (**Fig. 1a**, **Supplementary Table 1,2**). CTE diagnosis was performed by a neuropathologist and based on the presence of CTE pathognomonic p-tau lesions^10^ (**Fig. 1b**). Grey matter sulcus from the dorsolateral frontal cortex, one of the first brain regions affected in CTE, was processed for snRNAseq (**Fig. 1a**). After quality control and filtering, 170,717 nuclei of sufficient quality were clustered into 31 initial clusters and labeled based on expression of known cell type markers^11,12^ (**Fig 1c**, **Extended Data Fig 1l-n**). All major cell types were identified. Compositional analysis with scCODA demonstrated no significant differences in cell type abundance across pathological groups^13^ (**Fig 1d-f**, **Supplementary Fig. 1a**). Out of all major cell types, minimal RHI associated changes were observed in oligodendrocytes and oligodendrocytes precursor cells (**Supplementary Fig. 1b-i**), likely resulting from the grey matter focus of the current study. Thus we elected to focus further analyses on microglia, astrocytes, endothelial cells and neurons, consistent with prior studies^5,7,8,14^.

**Figure 1.**
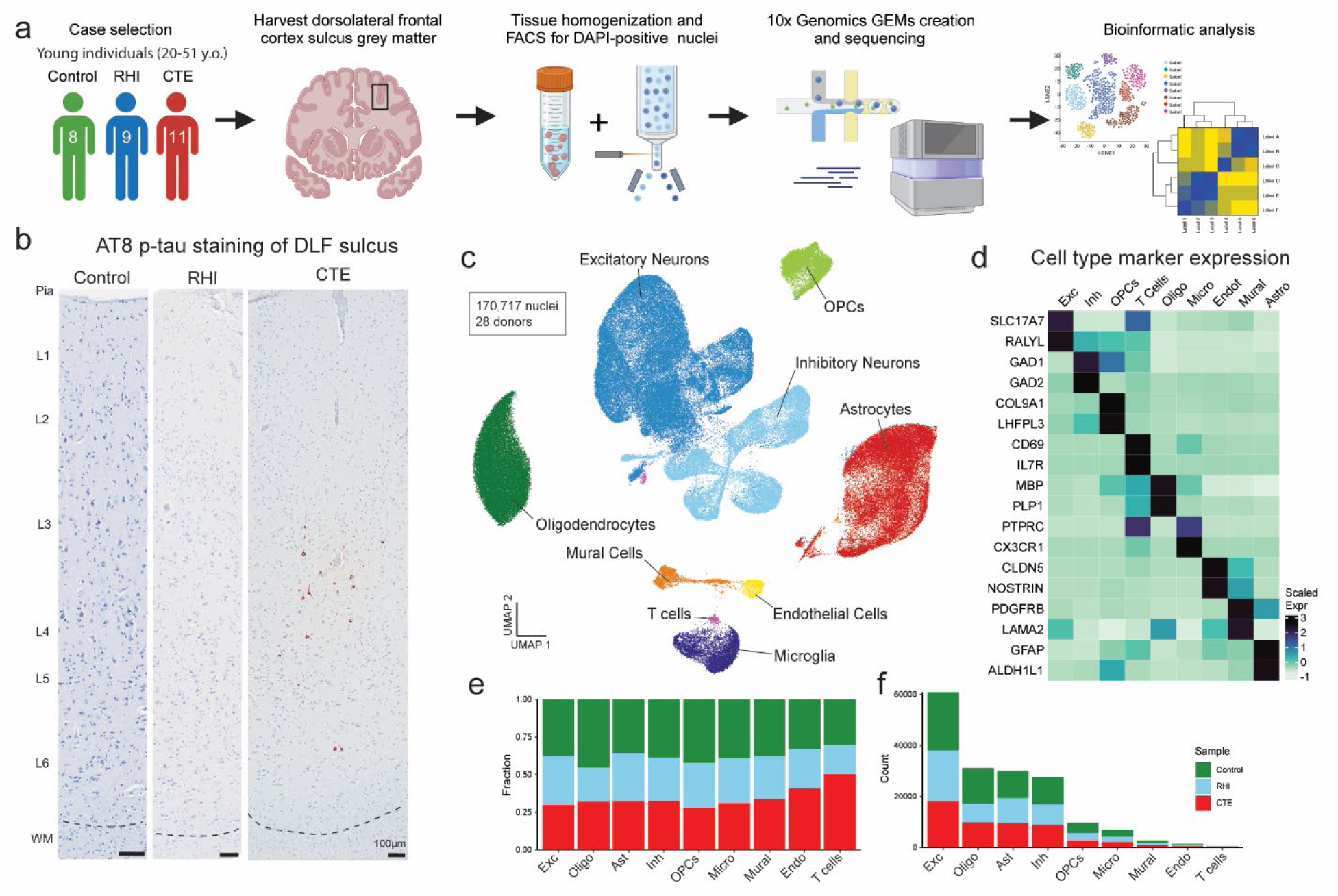
Cell type identification and cell proportion analysis across pathological groups. **a.** Diagram depicting experimental workflow. **b.** AT8 immunohistochemistry of dorsolateral frontal cortex depth of sulci, dashed line represents the grey-white matter interface. Scale bar, 100μm. **c.** UMAP of nuclei from all donors labelled for cell type based on cell-type marker expression. **d.** Expression of cell type markers across cell type clusters in (c). **e.** Stacked bar plot of pathological group fractions within cell type clusters. **f.** Stacked bar plot of cell type counts colored by pathological group.

### RHI exposure induces distinct microglial phenotypes

Based on previously demonstrated involvement of microglial inflammation in CTE and its important role in neurodegeneration, we examined microglial gene expression changes^5^. Analysis of 6863 microglial cells revealed eleven unique clusters (**Fig. 2a**). The microglia cluster size is consistent with other published studies and believed to be appropriately powered^11^. Cluster 10 contained 263 cells and expressed perivascular macrophage (PVM) genes CD163, F13A1, and LYVE1 and cluster 6 was composed of 108 cells expressed peripheral monocyte genes PTPRC, LYZ, and CR1 as previously observed ^11,15,16^(**Fig. 2b**).

**Figure 2.**
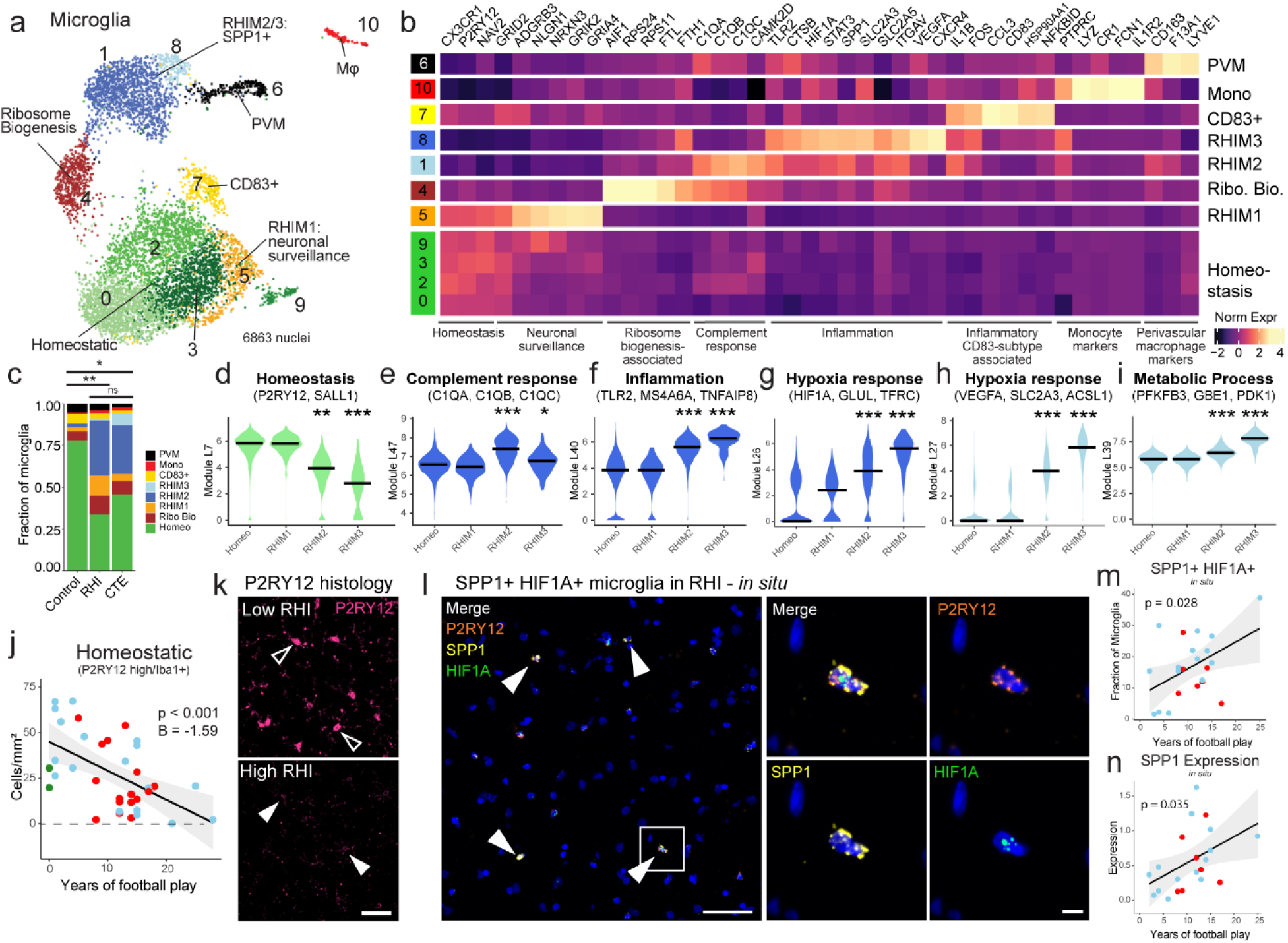
RHI Exposure induces distinct microglial phenotypes. **a.** UMAP of microglia colored by 11 Seurat clusters determined by unsupervised clustering. **b.** Heatmap of selected cluster DEGs annotated by function. **c.** Proportion of microglial subtypes per pathological group. Statistical analysis was performed using a chi-squared test. **d-i.** Violin plots representing the expression of Celda gene modules. Color represents the cellular subtype most associated with the module. Black line represents median statistic from ggsignif. Statistical analysis performed by linear mixed modeling correcting for patient-specific effects. *, p <0.05, **, p <0.01, ***, p.<0.001. **j.** Scatter plot depicting the density of immunohistochemically labeled homeostatic microglia (P2RY12 high /Iba1+) in the grey matter sulcus compared to years of football play, colored by pathological group identity. Statistical analysis performed by linear regression with age as a covariate. **k.** Representative image of P2RY12 immunofluorescent labeling (pink) in a low RHI and high RHI individual. Open arrows depict high P2RY12-expressing cells. Solid arrows depict low P2RY12-expressing cells. Scale bar 50μm. **l.** Representative image depicting *in situ* hybridization of SPP1+ (yellow)/HIF1A+ (green)/P2RY12+ (orange) microglia in an RHI-exposed individual. Solid arrows indicate triple-positive cells. White box indicates inset displayed on the right. Left scale bar 50μm, right scale bar 5μm. **m, n.** Scatter plot depicting SPP1+ HIF1A+ microglial fraction and microglial SPP1 expression in the grey matter sulcus compared to years of football play. Colored by pathological group status. Statistical analysis performed by linear regression.

Clusters 0, 2, 3, and 9 expressed classical microglial homeostatic genes CX3CR1, P2RY12, and NAV2 and were labelled as homeostatic microglia. Homeostatic clusters were significantly enriched for nuclei from control individuals compared to RHI controls and compared to CTE but not between RHI and CTE individuals (p = 0.048, 0.047, > 0.99, respectively, **Extended Data Fig. 2a**). Homeostatic microglial proportion decreased with increasing years of football play (p=0.004, β = -12.79). Cluster 7 highly expressed CD83, CCL3, and HSP90AA1, reminiscent of a possible pro-resolving phenotype recently identified in Alzheimer’s disease (AD)^17^. Cluster 4 had the highest differential gene expression of AIF1 (gene for Iba1) across clusters and was characterized by expression of FTL and FTH1 iron-associated genes along with expression of ribosomal-associated genes such as RPS24 and RPS11 (**Fig. 2b**).

The proportion of microglial subpopulations found in RHI and CTE individuals were significantly different from controls, with the emergence of Clusters 1, 5, and 8 in RHI and CTE individuals (**Fig. 2c**, **Extended Data Fig. 2b**). For simplicity, these clusters were labeled repetitive head impact microglia (RHIM) 1 through 3. Gene module analysis was performed with Celda to identify coexpression of possible cellular pathways across subclusters and linear mixed modeling statistical analysis was performed to compare gene module expression (**Fig 2d-i**, **Supplementary Fig. 2b**, **Supplementary Fig. 3**). Homeostasis-associated gene modules were significantly decreased in RHIM2 and RHIM3 (**Fig. 2d**).

Cluster 5, RHIM1, expressed neuronal-associated genes such as GRID2, GRIK2, and GRIA4, with top identified gene ontology (GO) terms including “synapse organization” (**Fig. 2b**, **Extended Data Fig. 2g**). Previous work has found that “satellite microglia” – microglia that closely contact neurons – increase in number following TBI and modulate neuronal firing activity^18^.

Cluster 1, RHIM2, were nearly evenly enriched for RHI and CTE (50% vs 46%, respectively) while cluster 8, RHIM3, were mostly CTE-enriched (83%). Transcriptionally, RHIM2 and RHIM3 were similar, displaying features of an inflammatory microglial phenotype with expression of SPP1, HIF1A, TLR2, IL1B, and CTSB (**Fig. 2b**, **Extended Data Fig. 2d**, **e**). SPP1 has been described as a general marker of inflammatory or activated microglia, potentially playing a role in synaptic engulfment in AD models^19^. SPP1 has also been described as an opsin for extracellular debris^20,21^ . GO analysis of RHIM2/3 DEGs identified “cytokine signaling in the immune system”, “positive regulation of immune response”, and “vesicle mediated transport” (**Extended Data Fig. 2g**). Gene module analysis demonstrated an increase in inflammation, hypoxia, and metabolic response in both RHIM2 and 3 compared to homeostasis clusters (**Fig. 2f-i**) providing orthogonal validation of GO and DEG analyses.

Some key differences were noted between RHIM2 and RHIM3. RHIM2 expressed C1QA, C1QB, C1QC, and CAMK2D the components and downstream effector of the C1q complement cascade known to drive aberrant synaptic engulfment in the neurodegenerative brain (**Fig. 2b**, **Extended Data Fig. 2f**)^22^. Gene module analysis further highlighted an increase in complement response in RHIM2 compared to homeostatic microglia (**Fig. 2e**). RHIM3 were characterized by upregulation of HIF1A and VEGFA, two central mediators of hypoxia, suggesting a potential response to or initiation of hypoxic conditions following RHI (**Extended Data Fig. 2f**). HIF1A also acts as a transcriptional regulator of numerous downstream inflammatory genes, and analysis of the transcriptional regulatory networks enriched in each cluster showed that RHIM3 expressed many genes regulated by HIF1A^23^ (**Extended Data Fig. 2h**).

To validate the reduction in the homeostatic microglial population, Iba1 and P2RY12 were co-immunolabelled and quantified in the sulcus of 35 individuals with 0 to 25 years of football play with or without CTE. Microglia were divided into high vs low expression P2RY12 expression. Homeostatic microglial densities (P2RY12 high/Iba1+) were significantly decreased with increasing years of football play (p < 0.001, **Fig. 2j, k**). Concurrently, non-homeostatic microglia (P2YR12 low/Iba1+) cells were positively correlated with increasing years of football play (p < 0.001, **Extended Data Fig. 2k**, **o**). Mirroring the snRNAseq results, CTE status was not significantly associated with homeostatic microglial densities when years of exposure were accounted for.

To verify the presence of RHIM2/3 cells and their relationship to pathology, *in situ* hybridization was performed to label microglia expressing RHIM2/3 marker genes SPP1 and HIF1A (**Fig. 2i, m**). P2RY12 was used as a marker for microglia as AIF1 (Iba1) is lowly expressed at the mRNA level evidenced by prior publication and the present snRNAseq data^23^. SPP1+/HIF1A+ microglia were quantified across 21 individuals with 2-25 years of football play with and without CTE (**Fig. 2l-n**). SPP1+/HIF1A+ microglia significantly increased with increasing years of football play in the cortical sulcus (p = 0.028, **Fig. 2m**).

There was no association between SPP1+/HIF1A+ microglia in the nearby cortical crest suggesting a regional specificity of this inflammatory phenotype (p = 0.53, **Extended Data Fig. 2l**). Analysis was performed to determine the layer specificity of SPP1+/HIF1A+ microglia, separating superficial and deep layers of the cortical sulcus. SPP1+/HIF1A+ microglia increased in both superficial layers 2-3 and deeper layers 4-6 (p = 0.039, 0.026, **Extended Data Fig. 2m**, **n**). This suggests that while the microglial inflammation is specific to the sulcus, there was no layer-wise specificity of this phenotype. Additionally, microglia increased expression of SPP1 with increasing years of football play (p = 0.035, **Fig. 2n**). CTE status and tau burden did not associate with the prevalence of SPP1+/HIF1A+ microglia (p = 0.34, 0.12, respectively).

Finally, we sought to compare our microglial populations to those described in published datasets, notably, Sun et al. published a dataset with over 100,000 microglia from over 400 individuals which was used as a comparison^23^. To do this we combined the datasets and reclustered them relative to one another and demonstrated good alignment of microglial subtypes (**Supplementary Fig. 4b**). Jaccard similarity scoring analysis confirmed alignment of RHIM2/3 with inflammatory, stress, phagocytic, and glycolysis-associated populations^23^ (**Extended Data Fig. 2i**,**j**).

Overall, these results suggest that RHI exposure induces an increase in neuronal surveillance and inflammatory microglial transcriptomic states before the onset of CTE. Inflammatory microglia are localized specifically at the sulcus in RHI-exposed individuals. These microglia may be involved in the initiation and maintenance of neuronal dysfunction, inflammation, and angiogenic processes present in CTE.

### Astrocytic responses to repetitive head impacts

Astrocytes play a key role in brain homeostasis in tasks such as neuronal and BBB maintenance and become reactive following RHI exposure and in neurodegenerative disease^14,24^. Four subtypes of astrocytes, Astro1-4, were identified based on stratification of pathological group identity, DEG analysis and gene module analysis (**Supplementary Fig. 5, Supplementary Fig. 6, Supplementary Table 19**). Although past work has suggested the importance of astrocytes in CTE, a limited astrocytic response was observed with only one subtype being enriched for individuals with RHI (Astro3). Astro3 upregulated genes and gene modules associated with astrocyte reactivity (CHI3L1, CD44, CLU, BCL6), inflammation (IL6R, IL1R1), and angiogenesis (HIF1A, NRP1, ANGPTL4, **Supplementary Fig. 5h**). These findings suggest that although there is pronounced astrogliosis associated with end stage CTE pathology, astrocytes might have a more subtle role in early disease.

### Endothelial angiogenic response to RHI

Next, due to the key involvement of vascular dysfunction in CTE, we characterized the vascular response to RHI exposure^8,9^ (**Extended Data Fig. 3a**). Known cell type markers and comparison to published dataset markers were used to identify 1762 endothelial cells, 913 pericytes, 487 fibroblasts, and 651 vascular smooth muscle cells^25^ (**Extended Data Fig. 3b**, c, e). Only fibroblasts displayed significant changes in total proportion across pathological groups, decreasing from controls to RHI and CTE and with loss associating with years of football play (p=0.048, 0.027, respectively, **Extended Data Fig. 3d**, **f**). Endothelial cells were further labelled for arterial, venous, and capillary cells through comparison of expressed genes to published datasets^25^ (**Extended Data Fig. 3c**, **e**). Capillary cells were then labeled Cap1-Cap4. Cap3 and Cap4, (Seurat Cluster 5 **Extended Data Fig. 3b**, **c**) displayed a slightly different transcriptomic profile with greater levels of collagen associated genes and showed overlap in expression of pericyte genes representing a potential transitional cell state but with greatest fidelity to endothelial cell expression (**Extended Data Fig. 3c**, **e**).

The proportion of endothelial cell subtypes differed significantly between RHI and control individuals and trended towards a difference between Control and CTE individuals (**Fig. 3a-b**). No difference was observed between RHI and CTE individuals (**Fig. 3b**). Two populations of capillary cells, Cap2 and Cap4 were enriched for RHI and CTE samples (p = 0.004, 0.005, **Extended Data Fig. 3h**). Cap2 cell fraction also increased with increasing years of football play (p = 0.014, **Extended Data Fig. 3j**). No differences were observed in total capillary cells in RHI and CTE compared to controls (**Extended Data Fig 3g**). Several canonical angiogenesis-associated genes such as HIF1A, ANGPT2, ANGPTL4, STAT3, CAMK2D, and NFKBID were significantly upregulated in Cap2 and Cap4 suggesting capillary cells in RHI-exposed groups may be responding to a local hypoxic environment (**Fig. 3c, d**). Three major complement regulatory proteins, CD59, CD55, and CD46, which inhibit complement-mediated cell lysis, were upregulated indicating a potential response to locally increased levels of complement (**Fig. 3c**). Vascular adhesion and transmigration-associated genes ICAM1, ICAM2, PECAM1, and CD99 were increased in Cap2 and Cap4, indicating an increased potential for monocyte, T cell, neutrophil, or other peripheral cell entry across the endothelium (**Fig. 3c**). Cap4 also displayed high expression of collagen genes (**Fig. 3c**). Module coexpression analysis using Celda was performed to identify co-expressed genes and possible cellular pathways across endothelial subsets. Statistical linear mixed modeling demonstrated that modules related to immune signaling, angiogenesis, response to growth factors and collagen associated modules were significantly upregulated in Cap2 and Cap4 subsets (**Fig. 3d**, **Supplementary Fig. 7, Supplementary Table 18**). GO analysis identified VEGFA signaling, cytokine signaling, and vasculature development as significantly upregulated terms in RHI-exposed endothelial cells (**Extended Data Fig. 3i**). We identified ITGAV as an endothelial gene that was significantly increased in Cap2 cells and increased in expression in RHI compared to control and CTE compared to RHI (**Fig. 3e**). To confirm its expression in the tissue we performed *in situ* hybridization paired with GLUT1 immunohistochemistry to label vessels and found an increase in the fraction of vessels expressing ITGAV with increasing years of football play (p = 0.027, **Fig 3. f, g**). Taken together, capillary cells undergo significant upregulation of angiogenesis and inflammation associated genes along with an increase in basement membrane components, identifying pathways that may underlie the known microvascular dysfunction after RHI and in CTE^8,9^.

**Figure 3.**
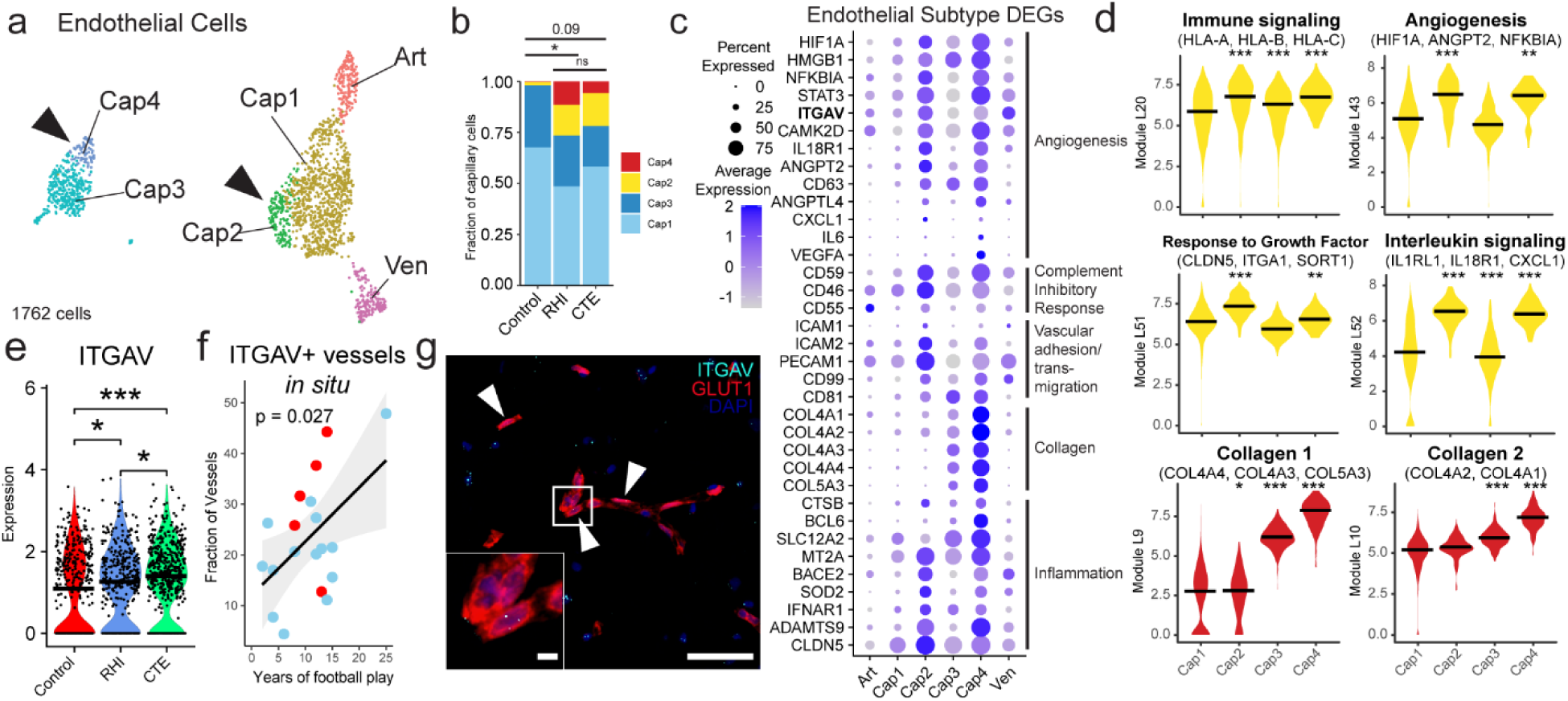
Endothelial angiogenic responses to RHI. **a.** UMAP of endothelial cells colored by endothelial cell subcluster. Solid arrows indicate RHI/CTE enriched clusters. **b.** Stacked bar plots of capillary subtype abundance across pathological groups. Statistical analysis performed using a chi squared test. **c.** Dot plot of selected upregulated RHI/CTE DEGs across endothelial subtypes annotated for function. **d.** Violin plots of Celda module expression across capillary subtypes. Black bars indicate median statistic from ggsignif. Statistical analysis performed with linear mixed effects model accounting for sample variability and comparing Cap2-4 to Cap1. **e.** Violin plot of ITGAV expression across pathological groups. Each dot representing a cell. Statistical analysis performed by Wilcoxon test from ggsignif. *, p <0.05, ***, p<0.001. **f.** Scatter plot of ITGAV+ vessel fraction in the grey matter sulcus compared to years of football play colored by pathological group status. Statistical analysis performed by linear regression. **g.** Representative image of ITGAV+ vessel. Solid arrows indicate ITGAV+ vessel. White box indicates inset. Left scale bar, 5μm, right scale bar 50μm.

### Synaptic transcriptomic changes and loss of sulcal excitatory cortical layer 2/3 neurons

Next, due to the known dysfunction and degeneration of neurons and synaptic dysfunction following head trauma and in neurodegenerative disease, we examined neurons, labeling subclusters using known layer-specific markers^12,26–31^ (**Fig. 4a**, **Supplementary Fig. 8e-j**). 47% of excitatory neuron DEGs were shared across RHI and CTE when compared to control and only 6% changed from RHI to CTE, suggesting that the greatest changes in excitatory neuronal transcriptional profiles occur with initial exposure to RHI (**Fig. 4b**). RHI and CTE gene expression was compared to controls and GO analysis of total neuronal population and layer-specific DEGs demonstrated that “modulation of chemical synapses” and “cell-cell adhesion” processes were enriched in both analyses (**Fig. 4c**, **Supplementary Fig. 9b**). Genes associated with synaptic transmission such as SYN3, SNAP91, NRG1, HSP12A1 the Hsp70 gene, and extracellular matrix binding proteins such as CNTN5, CLSTN2 were upregulated across several excitatory neuron layers. Inhibitory neuron layer-wise DEGs displayed 40% fewer DEGs than excitatory neurons with only 184 DEGs specific to RHI-exposed groups compared to controls. GO analysis of inhibitory neuron layer specific DEGs showed common upregulation of synapse associated genes such as SYN3 and SYN2 and across layers and downregulation of GABA receptor gene GABRA1 (**Supplementary Fig 9c**).

**Figure 4.**
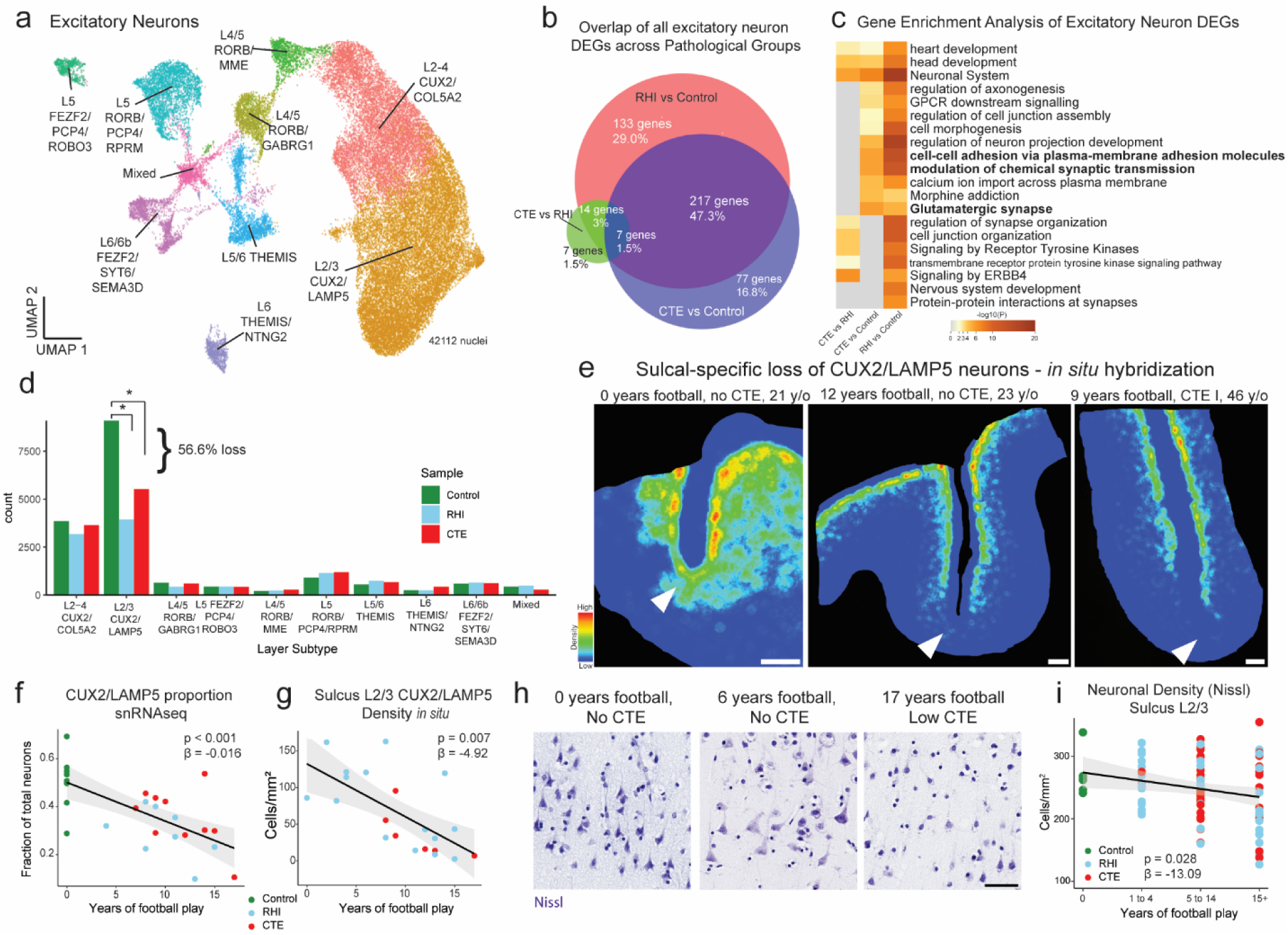
Synaptic transcriptomic changes and loss of sulcal excitatory layer 2/3 neurons. **a.** UMAP of excitatory neurons colored and labelled by layer subtype determined by expression of layer-specific markers. **b.** Venn Diagram depicting the overlap between DEGs from RHI vs Control, CTE vs Control, and RHI vs CTE comparisons. **c.** Heatmap of GO terms identified in comparisons listed in (b). **d.** Bar plot representing cell counts per pathological group for each excitatory neuron layer subtype. Statistical analysis performed by ordinary one-way ANOVA with Bonferroni correction. *, p<0.05. **e.** Representative density heatmap of CUX2/LAMP5 positive cells, solid arrows indicated depth of the cortical sulcus. Red indicates high cellular density; blue indicates low cellular density. Scale bar, 1 mm. **f.** Scatter plot showing the fraction of CUX2/LAMP5 neurons within total excitatory neurons in snRNAseq data against total years of football play colored by pathological group identity. Dots depict individual samples, line represents general linear model regression, grey shows 95% confidence interval **g.** Scatter plot showing cell density of CUX2/LAMP5 neurons in sulcal Layer 2/3 from *in situ* hybridization colored by pathological group identity compared to years of football play. Dots depict individual samples, line represents general linear model regression, grey shows 95% confidence interval. Statistics performed by general linear regression. **h.** Representative images of Nissl-stained neurons in superficial cortical layer 2/3, scale bar indicates 50μm. **i.** Scatter plot showing Nissl-stained neuronal densities across football exposure groups. Dots depict individual samples, line represents general linear model regression, grey shows 95% confidence interval.

Since neurodegenerative processes and head trauma exposure can be associated with neuronal loss, we investigated layer-specific cell composition in RHI and CTE individuals compared to controls. No pathological group enrichment was found in inhibitory neurons (**Supplementary Fig 9e, f**). However, differential abundance analysis of excitatory Layer 2/3 CUX2/LAMP5 neurons demonstrated a significant decrease in individuals with a history of RHI, regardless of CTE status (**Fig. 4d**, **Supplementary Fig. 9d**). These results were confirmed via multinomial dirichlet multinomial regression to account for the compositional nature of snRNAseq data^13^. RHI exposure individuals had an average of 56% fewer CUX2+/LAMP5+ neurons than age-matched unexposed controls. When measured by proportion of total neurons, loss of CUX2+/LAMP5+ neurons were also observed between RHI and CTE individuals and controls (p <0.01, <0.05, respectively **Supplementary Fig. 9d**). Neuronal loss was associated with the number of years of playing American football or, in the few cases with other types of contact sports play, total years of RHI exposure, independent of age at death (p<0.001, **Fig. 4f**, **Extended Data Fig. 4a**).

To determine the spatial localization of the neuronal loss and validate the snRNAseq results, quantitative histology with RNAScope *in situ* hybridization was performed using excitatory layer 2/3 neuron markers CUX2 and LAMP5. CUX2+/LAMP5+ neuronal density at the sulcus was negatively associated with years of football play (p = 0.007, β = -4.92) and highest level of football played (p = 0.033, β = -25.34, **Fig 4e, g**, **Extended Data Fig. 4b**). CUX2+/LAMP5+ cell density was significantly lower at the depth of the cortical sulcus compared to the nearby gyral crest, consistent with RHI specific damage and CTE pathology^32^ (**Fig. 4e**, **Extended Data Fig. 4d**,e). CUX2+/LAMP5+ cell densities at the crest were not associated with years of play, demonstrating a regional specificity of neuronal cell loss to the sulcus (p = 0.686, **Extended Data Fig. 4e**). CUX2+/LAMP5-(putatively CUX2+/COL5A2+) neurons are found intermixed with CUX2+/LAMP5+ neurons throughout layers 2-4 and are putatively exposed to similar levels of mechanical forces due to adjacent anatomical location. However, neuronal loss was observed to be specific to CUX2+/LAMP5+ expressing excitatory neurons across *in situ* and snRNAseq experiments, suggesting specific susceptibility of this population to RHI exposure (**Extended Data Fig. 4f**).

To further validate the association between years of football play and neuronal loss, total neuronal densities were determined using Nissl staining of 86 young individuals with 0-28 years of American football play. Individuals were grouped by 0, 1-4, 5-14, and 15+ years of football play based on previously defined thresholds for CTE risk^3^. Layer 2/3 sulcal neuronal density significantly decreased with increased binned years of football play independent of age at death (p = 0.028, β =-13.09, **Fig. 4h, i**). No association was found between years of football play and neuronal densities in deeper layers 4-6 or in layer 2/3 in the crest (p = 0.554, 0.571, **Extended Data Fig. 4g**, **h, k, l**).

As p-tau deposition has been shown to associate with neuronal loss in neurodegenerative disease, comparisons of neuronal densities to p-tau pathology in adjacent sections was performed. No association between neuronal loss and p-tau deposition was observed suggesting neuronal loss occurs prior to and independent of pathologic protein deposition in early stages of disease (p = 0.387 *in situ*, p = 0.825 Nissl, **Extended Data Fig. 4i**, j).

Microglia contribute and respond to neuronal loss^33^. To investigate potential relationships between the observed neuronal loss and loss of microglial homeostasis, layer-wise homeostatic microglial populations (P2RY12 high/Iba1+) from adjacent histological sections were compared to neuronal densities. Neuronal densities were significantly positively associated with homeostatic microglial populations in layers 2/3 (p = 0.047, B = 0.126, **Extended Data Fig 2p**). In contrast, in layer 4-6 neuronal densities were not associated with homeostatic microglia populations (p = 0.105, **Extended Data Fig. 2l**), suggesting loss of microglial homeostasis may be specifically localized to regions of neuronal loss.

Overall, these results show the first evidence that exposure to RHI alone may drive significant neuronal loss and dysfunction, which may help explain early symptom onset in young athletes without the presence of significant p-tau pathology. Additionally, the relationship between neuronal loss and loss of microglial homeostasis point to potential mechanisms of or responses to neuronal loss.

### Ligand-receptor pair analysis in RHI exposure and CTE

To determine signaling pathways that may be involved in the cellular response to RHI exposure and CTE pathology, ligand receptor (L-R) pair analysis was performed using multinichenet^34^. Two comparisons were run, RHI compared to control (**Fig. 5a** labeled “RHI”) to examine signaling occurring in the context of head trauma, and CTE compared to RHI (**Fig. 5a** labeled “CTE”) to investigate what signaling might be involved in the deposition of p-tau. In RHI-exposed individuals, microglial TGFB1 was identified as an important ligand, signaling to endothelial cells, astrocytes, neurons, and other microglia through TGFB1 receptors ITGAV, TGFBR2, TGFBR3, TGFBR1. SIGLEC9 and SPP1 were also found to be a major signaling hubs in RHI compared to control implicating the RHIM2/3 phenotype in RHI-associated signaling. In CTE compared to RHI, top microglial signaling pathways identified also included TGFB1 signaling and WNT2B and HLA-DRA signaling to astrocyte, microglia, endothelial cells and excitatory neurons. TGFB1 signaling has been previously implicated in the activation of neuroinflammation and induction of neuronal cell death in mild TBI^35^. Additionally, TGFB1 is involved with the fibrogenic response to mechanical stretch stimulus through ITGAV activation on endothelial cells and angiogenic responses through TGFBR2 signaling^36,37^.

**Figure 5.**
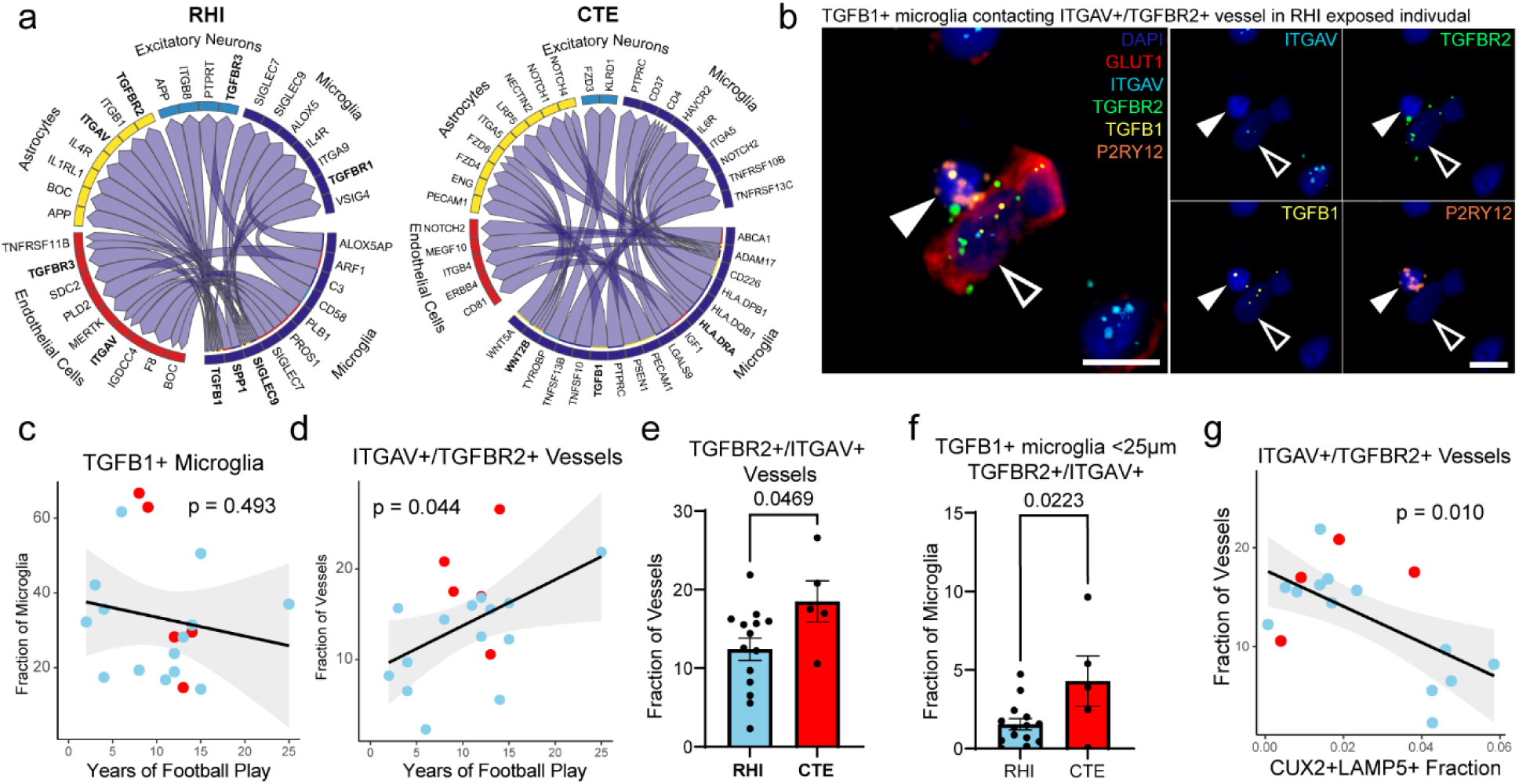
Ligand-receptor pair analysis in RHI exposure and CTE. **a.** Circos plots from multinichenet analysis depicting microglia as sender cells. RHI indicating RHI vs. Control contrast, CTE indicating CTE vs. RHI contrast. **b.** RNAScope *in situ* hybridization depicting a TGFB1+ (yellow) microglia (P2RY12+, orange, solid arrows) contacting a TGFBR2+/ITGAV+ (green, light blue) vessel (GLUT1, red, open arrows). Scale bars, 10μm. **c, d.** Scatter plots showing TGFB1+ microglia and ITGAV+/TGFBR2+ vessels in the grey matter sulcus compared to years of football play, color coded by pathological group. Statistical analysis performed by simple linear regression. **e.** Bar plot representing TGFBR2+/ITGAV+ vessels compared to CTE status, statistical analysis performed using a two-tailed t-test. **f.** Bar plot representing the proportion of TGFB1+ microglia within 25μm of a TGFBR2+/ITGAV+ vessel compared to CTE status, statistical analysis performed using a two-tailed t-test. **g.** Scatter plots depicting ITGAV+/TGFBR2+ vessels in the grey matter sulcus compared to the fraction of CUX2+/LAMP5+ neurons color coded by pathological group. Statistical analysis performed by simple linear regression.

In situ hybridization analysis was performed to label TGFB1 in microglia, and two of the receptors identified in endothelial cells L-R pair analysis: TGFBR2 and ITGAV (**Fig. 5b**). We hypothesized that TGFB1+ microglia would increase in proximity to ITGAV+/TGFBR2+ vessels to facilitate signaling. The prevalence of TGFB1 expressing microglia did not increase with the level of exposure nor with CTE status, however, an increase in ITGAV+/TGFBR2+ vessels was found with increasing years of football play and with CTE status (p = 0.493, 0.044, 0.047, respectively, **Fig. 5c, d, e**). Additionally, there was an increase in TGFB1+ microglia within 25μm of ITGAV+/TGFBR2+ vessels in CTE individuals compared to RHI exposed individuals without CTE (**Fig. 5f**). The increase in microglia-endothelial cell pairs is likely driven by an increase in endothelial ITGAV and TGFBR2 expression as opposed to an increase in TGFB1+ microglia, concurring with data from the microglia and endothelial sections. We then compared the prevalence of ITGAV+/TGFBR2+ vessels to CUX2+/LAMP5+ neuronal populations in adjacent sections to identify potential relationships between the identified signaling pathway and the observed neuronal loss. We found that with decreasing neuronal populations in the grey matter sulcus, there was an increase in ITGAV+/TGFBR2+ vessels (p = 0.010) (**Fig. 5g**). Overall, these findings identify a possible signaling pathway that may be implicated in the microglia-endothelial cell cross talk that may be implicated in the early pathological cascade of CTE pathology.

## Discussion

In this study we utilize a combination of single nucleus RNA sequencing, multiplex *in situ* hybridization, and immunohistological analyses to describe and validate a unique dataset of young individuals with exposure to RHI. We describe distinct microglial and endothelial subsets that emerge following RHI and persist with CTE, correlate with years of contact sport play, and associate with neuronal loss. Additionally, we observed a sulcus-specific loss of cortical layer 2/3 neurons that correlated with exposure to RHI prior to p-tau deposition. Finally, we identify a possible TGFB signaling cascade between microglia and endothelial that might drive early pathogenesis.

Microglial, astrocytic, and endothelial cell transcriptomic subtypes have been described in several neurodegenerative diseases and in severe traumatic brain injury, however this is the first study to demonstrate these changes in a young of a cohort with exposure to repeated non-concussive head impacts. Interestingly, hypoxia-associated changes are present across these three cell types, suggesting an important role for vascular dysfunction and bolstering previous evidence of vascular remodeling in CTE. Forces from head trauma disproportionately affect blood vessels, causing a lasting endothelial response affecting blood brain barrier integrity and oxygen delivery in affected regions^32^. Activated endothelium, local hypoxia, and a breached BBB may trigger a feedback loop activating astrocytes and microglia with each head impact. Repeated blows to the head in short succession likely reactivate an already inflamed system, disallowing sufficient time for full repair, and preventing a return to homeostasis. This is substantiated by the increased microglial activation observed decades after retirement from contact sports and found to correlate with the number of years of RHI exposure in this study and others^5^. Through this repetitive reactivation, the inflammatory response becomes self-sustaining and chronic, the mechanisms of which remain unclear. Some potential mechanisms identified in this study are increased collagen expression in endothelial cells potentially indicating an early endothelial fibrotic response. Identified through ligand-receptor analysis, TGFB1-ITGAV/TGFBR2 signaling between microglia and endothelial cells may represent a potential signaling pathway for the observed endothelial activation.

We observed a marked ∼56% decrease in superficial layer 2/3 excitatory neurons in RHI-exposed individuals at the depths of the cortical sulci, the region known to sustain the most mechanical force upon head trauma, and the initial region of p-tau accumulation in CTE^32^. This is the first study to demonstrate such a dramatic loss of a specific neuronal subtype in young individuals solely driven by RHI exposure. This is especially concerning considering several of the observed individuals had no neuropathologic protein deposition, suggesting neurodegeneration might begin sooner than CTE onset. Recent studies have demonstrated cortical thinning in frontal regions of high school football players, and cortical thinning and neuronal loss in postmortem individuals with CTE ^38,39^. Neuronal loss might explain symptoms of traumatic encephalopathy syndrome, the clinical criteria for antemortem CTE diagnosis, in young athletes^2,40,41^. Layer 2/3 neurons make cortico-cortical connections and, in the frontal cortex, are associated with depressive behaviors and moderation of stress^42^. Interestingly, layer 2/3 neurons have been shown to be vulnerable in other neurodegenerative and psychiatric disorders and are susceptible to p-tau accumulation in AD^12,27^. Therefore, one may speculate that superficial layer 2/3 excitatory neurons are highly susceptible to damage regardless of source and our data captured the loss across a range of RHI doses. Vulnerability has been hypothesized to be caused by the longer projections being more susceptible to trauma related diffuse axonal injury or the overall higher metabolic demand these cells need, however the exact mechanisms driving this susceptibility have yet to be elucidated. Although RHI damage is driving the early neuronal loss, it is likely that as p-tau deposition becomes more severe, neuronal death and dysfunction will become more related to pathogenic protein accumulation.

One limitation of our study is the small amount of tissue in each sample. CTE is an inherently patchy disease, and diagnosis is made based on the presence of a pathognomonic CTE lesion consisting of a focus of perivascular p-tau accumulation at the depth of the cortical sulcus. It is therefore possible that sampling may have missed regions of important cellular responses. Future studies of RHI-exposed individuals should aim to sample from several areas of the brain to improve detection of cellular responses. Additionally, due to the inherent difficulties in acquiring non-disease, non-RHI exposed young postmortem human samples, some control cases included in the Nissl quantification were female which may complicate direct comparisons to male athletes. However, no statistical correlation was found when sex was compared to neuronal densities.

These results highlight the growing concerns linked to long term RHI exposure from contact sports. The data presented here is some of the first direct evidence that demonstrates RHI-driven cellular perturbations occur prior to the development of CTE and can be observed in young individuals, many with no obvious brain pathology. Novel biomarkers and therapeutic interventions will be vital in identifying the early changes observed in contact sport athletes prior to developing neurodegeneration.

## Methods

### Neuropathological Diagnosis

Brain tissue was obtained from the CTE and the National Center for PTSD Brain Banks. Identical intake and tissue processing procedures occur with both brain banks. Four controls included in Nissl quantification were provided by the Iowa Neuropathology Resource Laboratory. Neuropathological examination was performed by board certified neuropathologists as described previously^10,43^. Diagnosis of CTE was determined using published consensus criteria^10,43^. Demographics such as athletic history, military history, traumatic brain injury history, and RHI history were queried during telephone interview with next of kin as detailed previously^10,43^. Institutional review board approval for brain donation was obtained through the Boston University Alzheimer’s Disease and CTE Center, Human Subjects Institutional Review Board of the Boston University School of Medicine, and VA Boston Healthcare System (Boston, MA). Individuals were included in the study based on frozen tissue availability, quality (RIN>4), and diagnosis. Exclusion criteria included neuropathological diagnosis other than CTE, moderate to severe traumatic brain injury directly prior to death, age of death greater than 51 or less than 25. Control cases did not have exposure to any RHI, were negative for any neurodegenerative disease, and did not carry any diagnosis of a neuropsychological disorder.

### Single Nucleus RNA Sequencing

Fresh frozen brain tissue was collected from the dorsolateral frontal cortex of each donor at the depth of the cortical sulcus. Visual delineation of grey/white matter was used to collect 50 mg of grey matter tissue. <u>Tissue was processed and cleaned of white matter prior to homogenization at two levels. First,</u> <u>when removing samples from frozen coronal slabs, the unbiased technician visually inspected and</u> <u>avoided white matter that could be adjacent to target grey matter. Second, immediately before tissue</u> <u>homogenization, a second technician inspects the tissue and removes any remaining white matter. This</u> <u>preparation allows for a highly specific grey matter enrichment</u>. Nuclei isolation and sorting were performed on two donor samples per day, randomizing for diagnosis and age. Tissue was kept on ice throughout nuclei isolation. Tissue was homogenized and lysed in NST Buffer with DAPI (146mM NaCl, 10mM Tris, 1mM CaCl2, 21mM MGCl2, 0.1%BSA, 0.1% NP-40, 40U/ml Protector RNase Inhibitor, DAPI) and snipped with scissors on ice for 10 minutes. Debris was removed using a 70μm filter. Cells were spun down and resuspended in nuclei storage buffer (2% BSA, 400U/mL Protector RNase Inhibitor) to reach a concentration of 500-1000 nuclei/μL. Nuclei were purified for DAPI positive cells with a FACS-Aria flow cytometer to remove debris and processed using the Chromium Next GEM Single Cell 3’ Reagents Kit V2 (10x Genomics) to create cDNA libraries. Samples were pooled in two batches sequenced with Azenta to a read depth of 30,000 reads/cell on an Illumina NovaSeq.

### Processing, Quality Control, and Clustering of Single Nucleus RNA Sequencing Data

CellRanger v6.0.1 was used to align reads to the GRCH38 reference and generate filtered count matrices containing 233,555 across all samples. The “runCellQC” function in the *singleCellTK* R package was used to generate quality control metrics and doublet calls^44,45^. Contamination from ambient RNA was identified using decontx using the full raw matrix as the “background” for each sample^46^. Nuclei were removed if they had ambient RNA contamination fraction greater than 0.3, mitochondrial or ribosomal percentage greater than 5%, total counts less than 750, or genes detected less than 500. The data was not down sampled to maximize capture of rare populations. The Seurat workflow within the *singleCellTK* package was used for clustering starting with the decontaminated counts from decontx^47^. Briefly, the data was normalized and scaled using runSeuratNormalizeData and runSeuratScaleData. Highly variable genes were identified using runSeuratFindHVG with method ‘vst’. Principle components were determined using runSeuratPCA. UMAP dimensionality reduction was calculated using runSeuratUMAP. Clusters across all cell types were identified using the runSeuratFindClusters function at a resolution of 0.3. After initial clustering all the cells, clusters that were predominantly doublets (>50%) were removed and produced the final dataset of 170,717 nuclei (Extended Data Fig. 1h-k). Associations with post-mortem interval (PMI), age at death, and sequencing batch were performed using Pearson’s correlation analysis in R (**Supplementary Fig. 4a**). Age at death was associated with only excitatory neuron L5_FEZF2_PCP4_RPRM and inhibitory neuron PVALB_SCUBE_PTPRK proportions. Therefore, age was not included in regressions performed with sequencing data. PMI correlated with only one microglial subtype (RHIM1), perivascular macrophages, an excitatory neuron subtype (L2_4CUX2_COL5A2) and several oligodendrocyte subtypes. Sequencing batch was associated with one cluster of OPCs and was therefore not included in analyses.

All GO analysis was performed using MetaScape default settings^48^. DEG lists for all comparisons available in Supplementary Tables 6-16.

### Cell type identification

Cell type markers verified by previous human snRNAseq studies were used to identify clusters that belonged to individual cell types (**Extended Data Fig. 1m**, n). Cell types were subsetted out using subsetSCEColData and reclustered by the same Seurat method described above with the addition of running Harmony to account for sample-to-sample variability^49^. Clusters expressing high levels of >1 cell type marker were removed. Excitatory and inhibitory neurons identified from the full dataset were clustered together to determine neuronal subtypes. Four clusters (1, 2, 19, 21) were found to express low levels of neuronal genes and astrocytic genes (SLC1A2, SLC1A3), and were single-batch enriched (80-90%) therefore these clusters were not included in downstream analysis (**Supplementary Fig. 8a-d**).

### Celda Module Analysis

Gene co-expression modules were identified using Celda^50^. Celda utilized Bayesian hierarchical linear mixed effects models to identify modules of genes that are expressed together. A workflow overview can be found in **Supplementary Figure 2**. Celda was run on cellular subtypes to determine module scores on a cell-wise basis and plotted across cellular subtypes. Statistical analysis of module enrichment was performed using a linear mixed effects model using sample ID as a covariate. For microglia, cell subtypes were compared to homeostatic microglia as a baseline, for endothelial cells Cap1 was used, for astrocytes Astro1 (homeostatic astrocytes) were used as a baseline. Module genes and statistical analysis can be viewed in **Supplementary Tables 17-19**, module expression across cell subtypes can be viewed in **Supplementary Figures 3, 6, 7**, analysis code is available on GitHub.

### Multinichenet

Ligand-receptor pair analysis was performed using multinichenet, an adaptation of nichenet that allows for comparison across more than two condition groups. Briefly, this method uses known datasets of ligand-receptor pairs and their downstream targets to identify potentially upregulated cell signaling pathways across cell types accounting for differential expression of genes across groups. Multinichenet also uses prioritization of top ligand receptor pairs to help identify signaling pathways of interest. Contrasts for differential gene expression were set as RHI versus Control, and CTE versus RHI to determine RHI and CTE-specific signaling pathways. Finalized cell type objects were combined and run through the multinichenet pipeline with the exclusion of T cells due to low cell numbers. Analysis was performed without alteration to publicly available code, save for the contrasts used.

### Histological Tissue Processing

Formalin fixed, paraffin embedded tissue was sectioned and labelled as previously described^51^. Briefly, 10μm sections were allowed to dry, baked, dewaxed, and rehydrated prior to antibody labelling. For immunofluorescent staining, epitope retrieval was performed using a pH 6 or pH 9 buffer and boiling for 15 minutes in the microwave. Sections were blocked for 30 minutes at room temperature with 3% donkey serum and primary antibodies (**Supplementary Table 4**) were conjugated for 1 hour at room temperature. Secondary antibodies were conjugated for 30 minutes, and Opal TSA dyes were incubated for 10 minutes. Slides were coverslipped with ProLong Gold Antifade mounting medium (Invitrogen) and imaged at 20x or 40x on a Vectra Polaris whole-slide scanner with the appropriate filters. Images were spectrally unmixed using inForm software prior to image analysis. For Nissl staining, sections were hydrated and stained in 0.01% thionin for 20-40 seconds and dehydrated back to xylene before coverslipping in Permount mounting media and imaging on an Aperio GT450 scanner at 40x.

### Single molecule fluorescent mRNA in situ hybridization and IHC codetection

Tissue was embedded in Optimal Cutting Temperature medium (Sakura Tissue-Tek) and was brought to cryostat temperature (−20⁰ C) before cutting. Chuck temperature was raised to -12⁰/ -10⁰C for optimal cutting conditions. Tissue was sectioned at 16 µm thickness onto Fisher SuperFrost slides. Direction of tissue orientation relative to the depth of the cortical sulcus was randomized across samples. Sections were fixed in cold 4⁰C 10% Neutral Buffered Formalin for 60 minutes and dehydrated in 50%, 70%, 100%, and 100% ethanol for 5 minutes each at room temperature. Fluorescent *in situ* hybridization was performed using RNAScope kits (Advanced Cell Diagnostics) optimized on the Leica BOND Rx automated slide staining system. Slides were pretreated with protease for 15 minutes. Opal TSA dyes were used for visualization at a concentration of 1:300-500. A positive and negative control probe was run for each block before staining with targeted probes. For immunohistochemical codetection of p-tau and GLUT1, sections were run through the RNAScope protocol as described and then manually stained with the AT8 or GLUT1 antibody (**Supplementary Table 4)** with the immunohistochemical protocol described in the Histological Analysis section less the antigen retrieval.

### Image Analysis

Analysis of fluorescent RNAScope *in situ* hybridization (FISH) was performed in Indica Labs HALO using the FISH v3.2.3 algorithm or the FISH-IF v2.2.5 algorithm. Thresholds for FISH probe positivity for was set manually for each probe (HIF1A, SPP1, P2RY12, ITGAV, TGFB1, TGFBR2, LAMP5, CUX2) and kept consistent across samples. It should be noted that SPP1 is not exclusively expressed by microglia, and DEG analysis demonstrated that only oligodendrocytes showed elevated expression of SPP1 in our dataset (**Supplementary Table 6b**). However, colocalization with microglia markers allows for a microglia specific count of SPP1 activity. Gene expression was determined by the “Probe Cell Intensity” in HALO because this measure is agnostic to manual single copy intensity settings. The background on GLUT1 staining in FISH sections was variable due to protease treatment from RNAScope and thresholds were manually adjusted to remove background staining. Vessel proximity analysis was performed by evaluating TGFB1+/P2RY12+ cells and GLUT1+/TGFBR2+/ITGAV+ cells and using the Proximity Analysis algorithm in the HALO spatial analysis settings. The number of unique marker-positive microglia/vessel pairs within 25µm were evaluated. Density heatmaps for CUX2/LAMP5 staining were created using the Density Heatmap function within HALO Spatial Analysis. Depiction of how the sulcus and crest were annotated can be found in **Extended Data Fig. 4d**. To validate consistency between image analyses methods and snRNAseq results, seven samples that were included in both RNA scope and snRNAseq methods were compared and cellular proportions of CUX2+/LAMP5+ neurons significantly correlated (p =0.02, **Extended Data Fig. 4c**).

Analysis of IHC protein staining was performed using the HALO Object Colocalization v2.1.4 and HighPlex v4.3.2 algorithm. Microglial P2RY12 was assessed by DAPI+ Iba1+ nuclei and P2RY12 high/low thresholds were set manually. High P2RY12 was defined as having at least 70% of the nucleus stained, low P2RY12 was defined as less than 70% of the nucleus stained as demonstrated visually in **Fig. 2f**. Only 5.4% of all Iba1+ or P2RY12+ cells were P2RY12+/Iba1-suggesting that 94.6% of labelled microglia were assessed. Iba1+ P2RY12-cells may have been captured in our P2RY12-low categorization however previous studies have shown that these cells are low in abundance and likely represent infiltrating macrophages which have been shown to be present mainly at lesioned vessels in CTE which are also sparse in our cohort^52,53^.

Analysis of Nissl staining was performed using the HALO Nuclei Segmentation AI algorithm. Neurons were selected for training based on previously published criteria^54^. Briefly, the classifier was given examples of brain parenchyma annotated for neurons which were considered cells with a Nissl-positive cytoplasm and a visible nucleus **(Supplementary** Fig. 9h). Nissl+ densities across batches were not significantly different and statistical tests of Nissl densities were corrected for staining batch. For FISH and Nissl sections, the depth of the cortical sulcus was defined and annotated as the bottom third of a gyral crest and sulcus pair. Layer 2/3 and layers 4-6 were annotated using layer-specific FISH markers or for Nissl by an expert observer.

### Inclusion & Ethics Statement

The research has included local researchers through the research process and is locally relevant with collaborators. All roles and responsibilities were agreed amongst collaborators ahead of the research. The research was not severely restricted in the setting of researchers. The study was approved by the Institutional review board through the Boston University Alzheimer’s Disease and CTE Center, Human Subjects Institutional Review Board of the Boston University School of Medicine, VA Bedford Healthcare System, VA Boston Healthcare System, and Iowa Neuropathology Resource Laboratory. The research did not result in stigmatization, incrimination, discrimination, or risk to donors or research staff. No materials have been transferred out of the country. Local and regional research relevant to the study has been included in the citations.

## Statistics

Analyses were performed using GraphPad Prism 10, SPSS v.29, and R packages ggsignif, muscat, scater, and Python package scCoda. Dirichlet multinomial regression was used to test for cell type and excitatory neuron cell type enrichment using the scCoda v0.1.9 Python package^13^. Celda module expression was evaluated using linear mixed effects modeling accounting for individual sample differences. Comparisons of cell type proportions across the three pathological groups were performed using ANOVA with Bonferroni correction, Brown Forsyth with Dunnett post-hoc test, or chi-squared test as indicated in figure legends. Comparison across control and RHI-exposed groups was performed with a t-test with Welch correction or Mann Whitney U test, as indicated in figure legends. Evaluation of *in situ* hybridization analysis was performed using linear regression. P-tau burden was normalized using log10 transformation of positive area density. Nissl+ neuron count comparisons to years of exposure was assessed using linear regression and correcting for age at death and staining batch. Jaccard similarity scoring was performed using the GeneOverlap package by comparing lists of DEGs. All DEGs were filtered by a log2fc of 0.15 and FDR of <0.05. Chi-squared tests for cellular abundance were performed using the base R chisq.tst function. Gene ontology analysis p values were acquired through MetaScape analysis. Years of football play was used as a variable for exposure throughout the text instead of total years of play (which includes exposure from all sports) played because it was a more consistent predictor of cellular changes.

## Supporting information

Supplementary Table of Contents

## Acknowledgements

We would like to extend our gratitude to the brain donors and their families without whom this work would be impossible. We thank the clinical and neuropathology teams at the BU CTE Center who perform clinical interviews with next of kin and routine tissue processing and Dr. Doug Rosene, Bryce Conner, and Sarah Horowitz for methodological support.

## Extended Data Figures

**Extended Data Figure 1.**
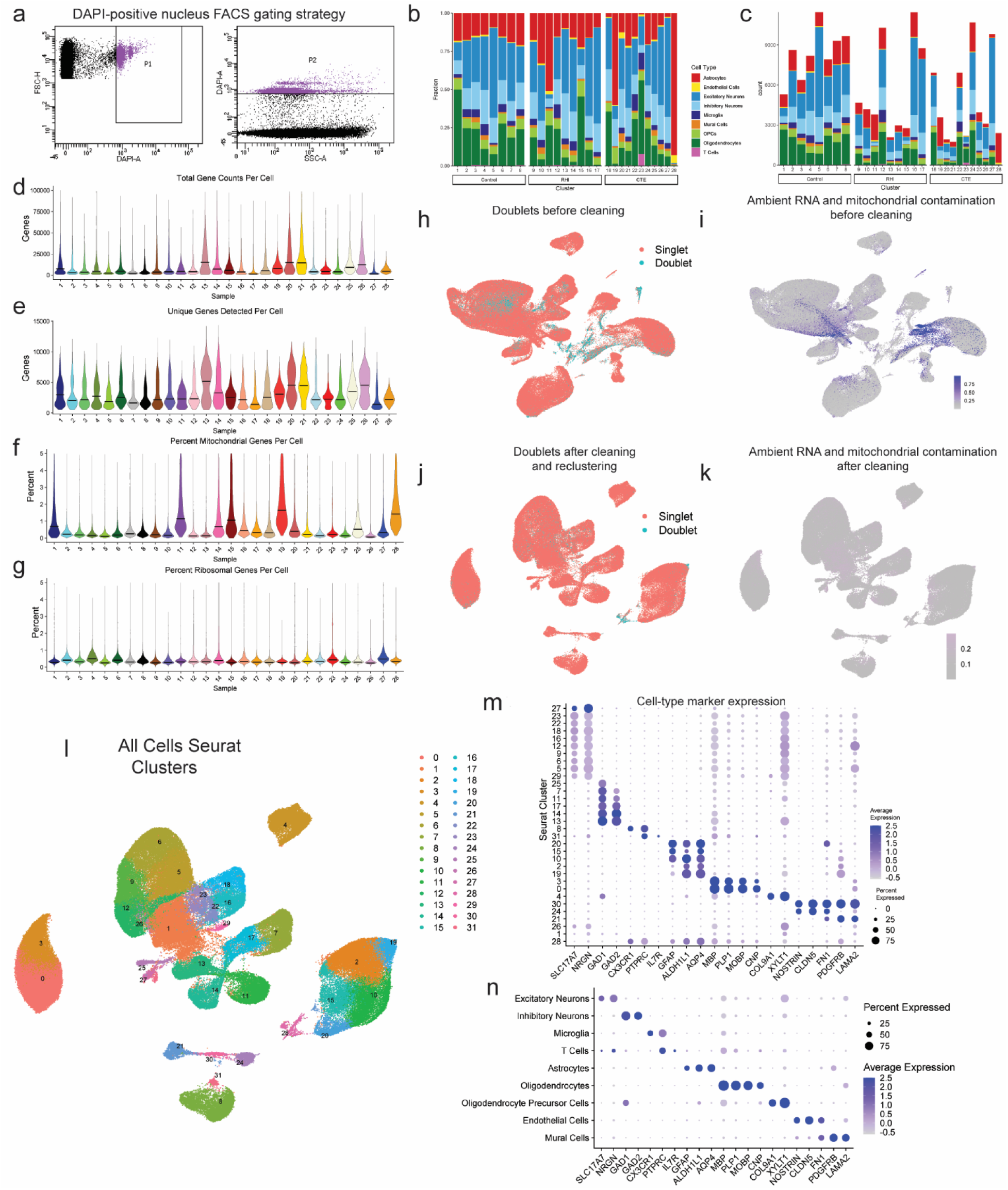
Dataset quality control and cell type marker validation. **a.** Fluorescence activated cell sorting gating strategy of DAPI-positive nuclei. **b.** Stacked bar plot representing the proportion of cell type per donor. **c.** Stacked bar plot representing the cell type counts per donor. **d-e.** Violin plots for each donor of **(d)** total gene counts per cell, **(e)** unique genes detected per cell, **(f)** percent of mitochondrial genes detected per cell, and **(g)** percent ribosomal genes detected per cell. Line represents median. **h, i** UMAP of full dataset before cleaning colored by **(h)** doublet or singlets or **(i)** mitochondrial contamination. **j, k.** UMAP of full dataset after cleaning colored by **(j)** doublets or singlets or **(k)** mitochondrial contamination. **l.** UMAP of full dataset colored by Seurat clusters. **m.** Dot plot of cell type marker expression across Seurat clusters depicted in **(l)**. **n.** Dot plot of cell type marker expression in annotated cell type clusters.

**Extended Data Figure 2.**
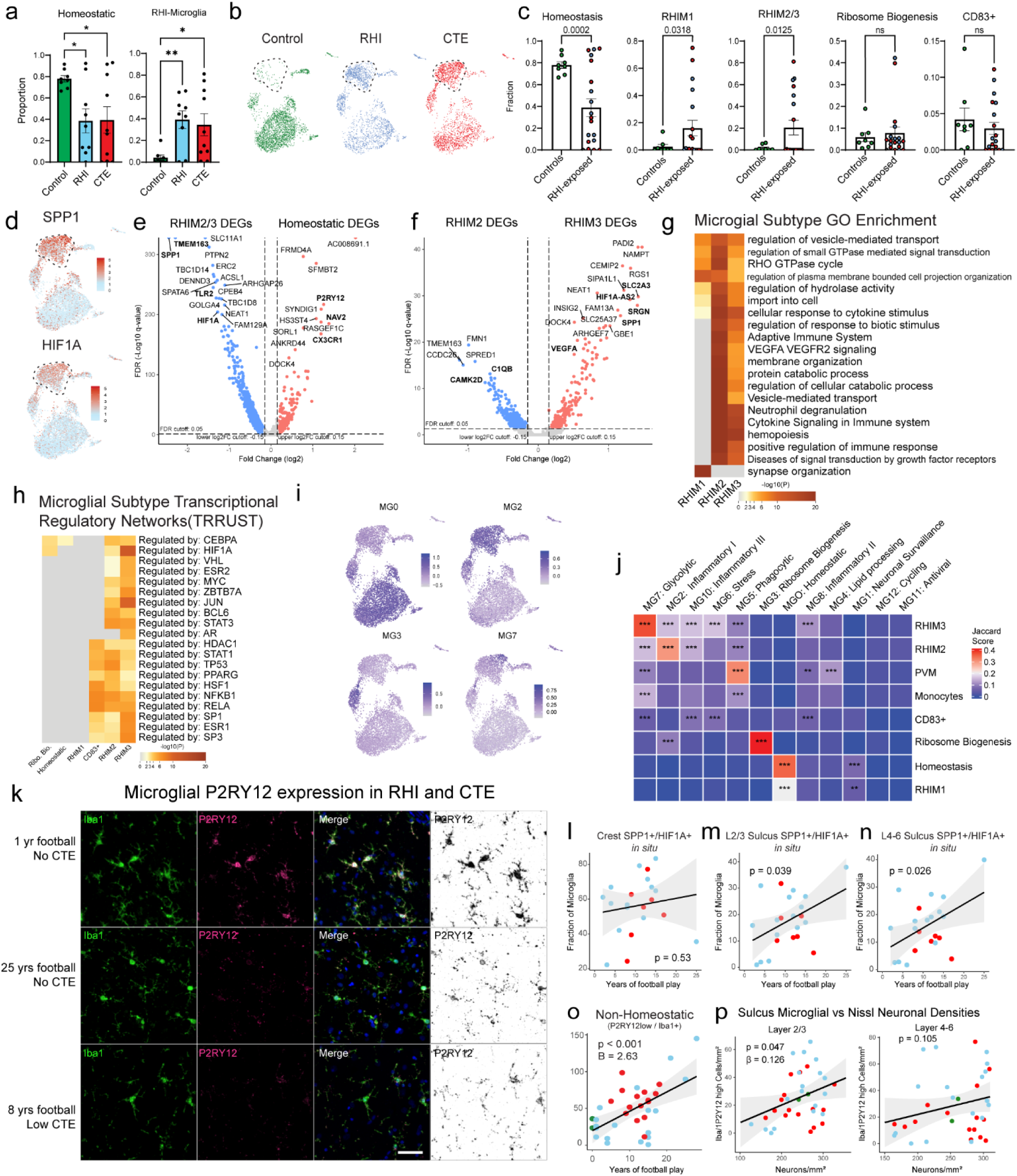
Microglial cluster GO analysis, histology, and validation. **a.** Bar plots showing fraction of homeostatic and RHI microglia across pathological groups. Statistics performed by ANOVA with Bonferroni correction. *, p < 0.05, **, p <0.01. **b.** UMAPs showing microglia from each pathological group. Dashed line highlighting RHIM2/3. **c.** Bar plots showing microglial subtypes across control and RHI-exposed individuals (RHI and CTE). Statistical analysis performed by two-tailed t-test or Mann Whitney U test with Welch correction. **d.** UMAPs showing microglial expression of SPP1 and HIF1A. Dashed lines indicate RHIM2/3. **e, f.** Volcano plots showing DEGs between RHIM2/3 and homeostatic microglia (e) and RHIM2 and RHIM3 (f). **g.** Heatmap of GO analysis of RHI microglia. **h.** Heatmap of transcriptional regulatory network analysis of microglial subtype DEGs. **i.** UMAPs depicting microglia colored for module scores of microglial subtypes from Sun et al. **j.** Heatmap depicting Jaccard score similarity analysis between Sun et al. and current study microglial DEGs. **, p<0.01, ***, p<0.001. Statistical analysis performed using GeneOverlap package and Jaccard analysis settings. **k.** Representative images of P2RY12(pink/black), Iba1 (green) immunofluorescent labelling in a low RHI, high RHI, and CTE individual. P2RY12 was also provided in an inverted pseudo black/white scale to better visualize expression since it can be present, but weakly expressed and sometimes difficult to observe. Scale bar, 50μm**. l, m, n.** Scatter plots depicting SPP1+/HIF1A+ microglial fraction in the grey matter (l) crest, (m) L2/3 Sulcus (n) layers 4-6 sulcus colored by pathological group status compared to years of football play. Statistical analysis performed by linear regression. **o.** Scatter plot depicting P2RY12 low/Iba1+ microglial densities in the grey matter sulcus compared to years of football play. Statistical analysis performed by linear regression with age included as a covariate. **p.** Scatter plot comparing homeostatic microglial densities to Nissl+ neuronal densities in layers 2/3 (left) and layers 4-6 (right). Statistical analysis performed by linear regression with age as a covariate.

**Extended Data Figure 3.**
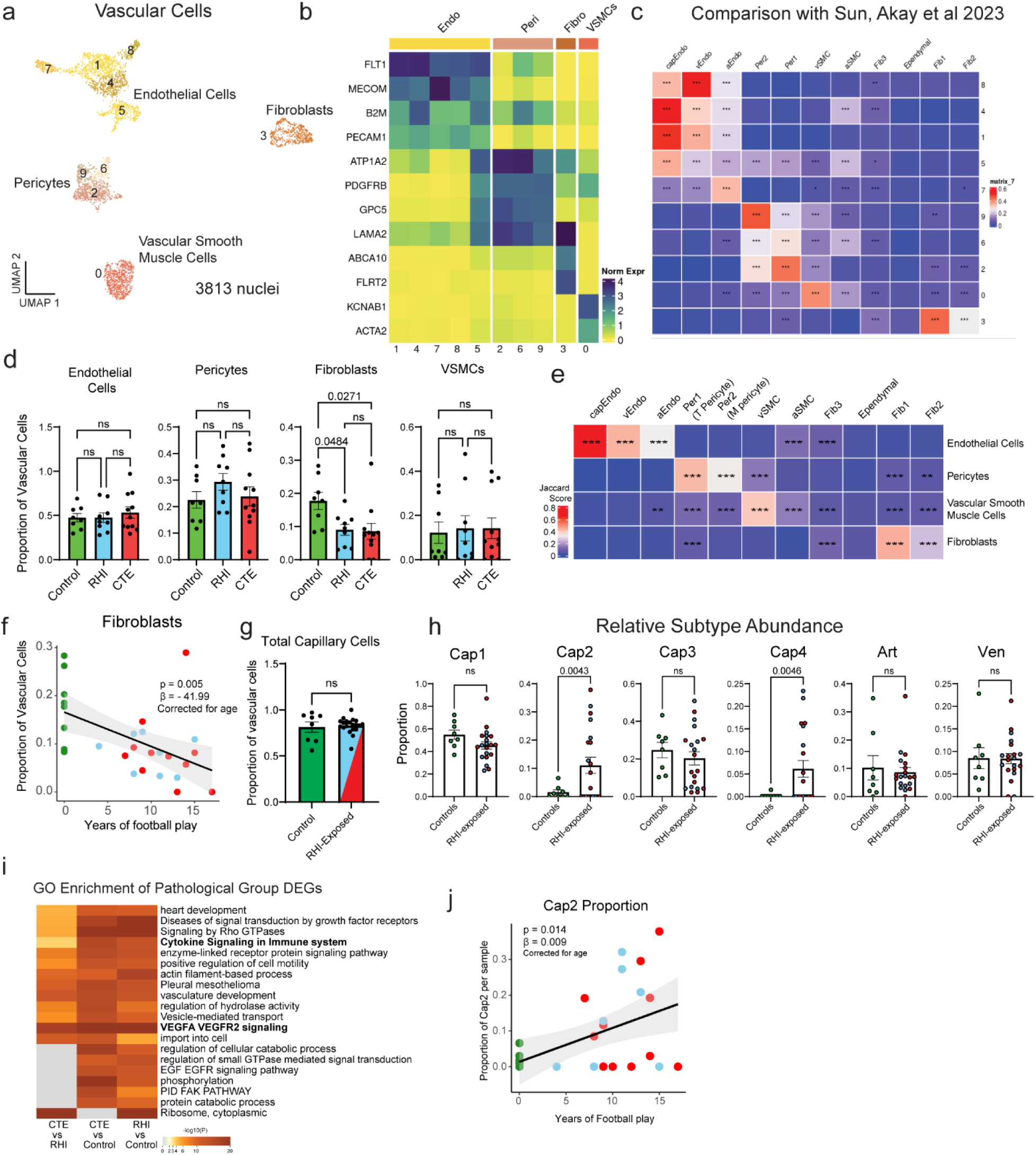
Vascular cell subtype identification and proportion analysis. **a.** UMAP showing all vascular cells colored by Seurat clustering. **b.** Heatmap depicting vascular cell marker expression. **c.** Heatmap depicting Jaccard scoring of vascular cell Seurat cluster DEGs compared to Sun and Akay et al. vascular subtype DEGs **, p<0.01, ***, p<0.001. **d.** Bar plots depicting pathological group proportions of vascular subtypes, bar represents mean, error bar represents standard error of the mean, dots represent individual samples. Statistical analysis performed by ANOVA with Bonferroni correction. **e.** Heatmap depicting Jaccard scoring of vascular cell subtype DEGs compared to Sun and Akay et al. vascular subtype DEGs **, p<0.01, ***, p<0.001. **f.** Scatter plot of fibroblast proportion or Cap2 proportion compared to years of football play from snRNAseq dataset, colored by pathological group status. Statistical analysis performed by linear regression with age as a covariate. **g, h.** Bar plots of total capillary and relative endothelial cell subtype distribution across control and RHI-exposed samples, dots represent individual donors and are colored by pathological group identity. Bar indicates mean, error bars indicate standard error of the mean. Statistical analysis was performed by two sided Mann-Whitney U test. **i.** GO enrichment analysis of DEGs from depicted comparisons.

**Extended Data Fig. 4.**
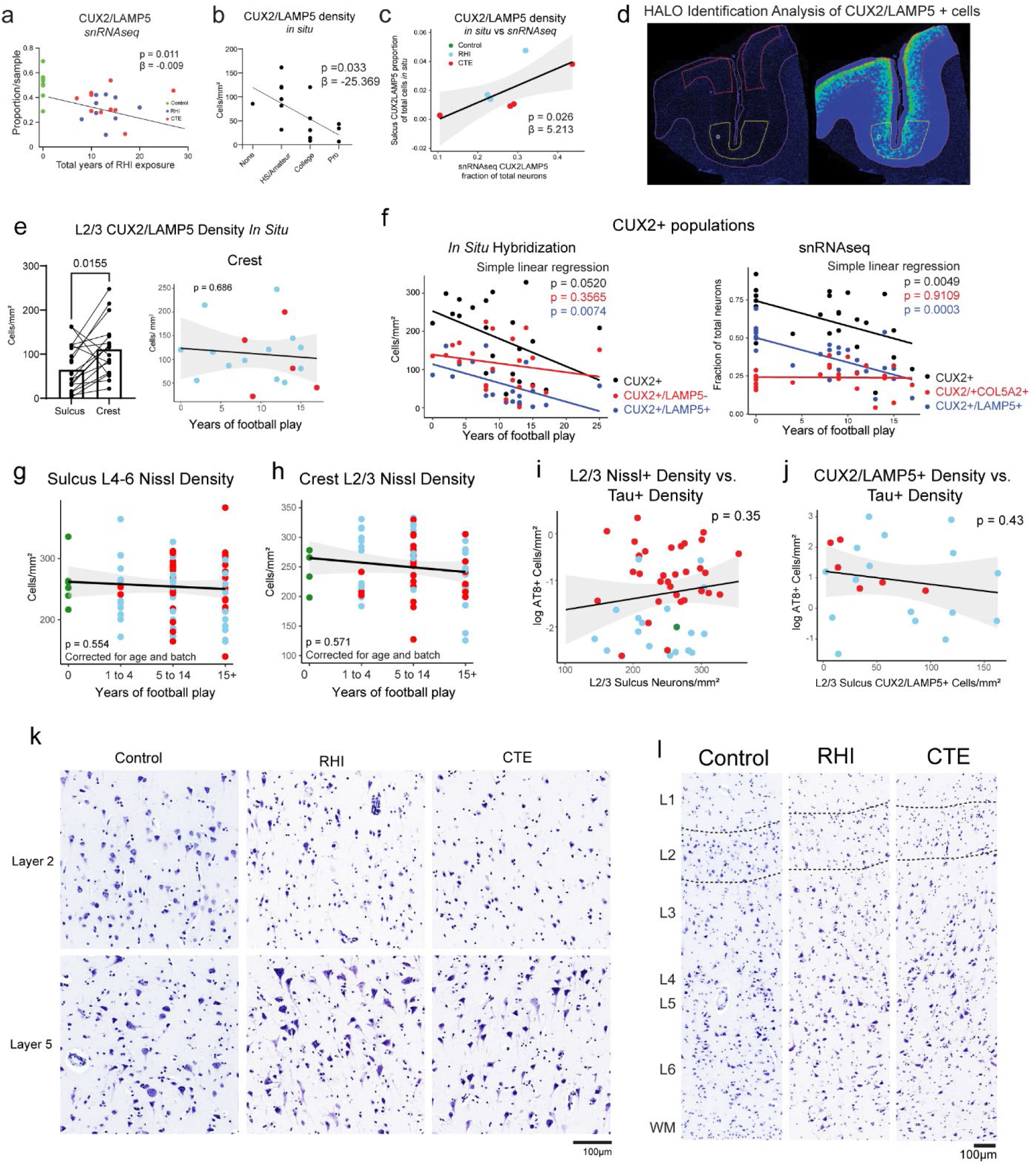
Layer 2/3 neurons are selectively lost in the grey matter sulcus and do not associate with tau pathology. **a.** Scatter plot of CUX2/LAMP5 proportion from snRNAseq against total years of football play colored by pathological group. Statistical analysis performed by linear regression, depicted as line. **b.** Scatter plot showing CUX2/LAMP5 density from *in situ* hybridization compared to highest level of football played. Statistical analysis performed by simple linear regression. Dots represent individual samples; line shows linear regression. **c.** Scatter plot of CUX2/LAMP5 cells identified by *in situ* experiment compared to proportion of CUX2/LAMP5 neurons from snRNAseq experiment. Statistical analysis performed by simple linear regression, depicted as line with 95% confidence intervals in grey. **d.** Representative image showing the annotation of sulcus (yellow line) and crest (red line) layer 2/3. **e.** (left)Bar plot depicting layer 2/3 CUX2+/LAMP5+ neuronal density in the sulcus and crest. Statistical analysis performed by paired t-test. (right) Scatter plot showing layer 2/3 CUX2+/LAMP5+ neuronal density in the crest compared to years of football play, colored by pathological group status. Statistical analysis performed by simple linear regression. **f.** Scatter plot depicting all total CUX2 populations and subpopulations in snRNAseq and *in situ* hybridization experiments compared to years of football play. Statistical analysis performed by linear regression. **g, h, i, j.** Scatter plots depicting (g) Sulcus layers 4-6 Nissl+ density compared to binned years of football play, (h) Crest layer 2/3 Nissl+ density compared to binned years of football play, (i) L2/3 Nissl+ density compared to log tau+ density, (j) CUX2+/LAMP5+ density *in situ* compared to log tau+ density, Colored by pathological group status. Statistical analysis performed by simple linear regression, (g, h) corrected for age and staining batch. **k, l.** Representative images of Nissl staining across cortical layers depicting neuronal loss in superficial layers in RHI and CTE individuals. Scale bars, 100μm.

## Funding

This work was supported by grant funding from: NINDS (F31NS132407), NIH (U19-AG068753), NIA (AG057902, AG062348), NINDS (U54NS115266), National Institute of Aging Boston University AD Center (P30AG072978); Department of Veterans Affairs Biorepository (BX002466), and the Department of Veterans Affairs Career Development Award (BX004349), BLRD Merit Award (I01BX005933). The views, opinions, and/or findings contained in this article are those of the authors and should not be construed as an official Veterans Affairs or Department of Defense position, policy, or decision, unless so designated by other official documentation. Funders did not have a role in the design and conduct of the study; collection, management, analysis, and interpretation of the data; preparation, review, or approval of the manuscript; or decision to submit the manuscript for publication.

## Competing Interests

The authors report no conflicts of interest.

## Author Contribution

Article conceptualization and writing performed by MLB and JDC. Experiments were performed by MLB, PY, KB, SC. Analysis was performed by MLB, NP, YW, JDC. Computational support provided by SM, JC. Neuropathologic diagnosis performed by MH, VEA, BH, TDS, ACM.

## Supplementary Figure Legends

**Supplementary Figure 1. Cell type proportions, OPCs, and Oligodendrocytes. a.** Bar plots of overall cell type proportions across pathological groups with each dot representing a sample, bars represent the mean, error bars represent standard error of the mean. Statistical analysis performed by ANOVA with Bonferroni correction. **b.** UMAP depicting OPCs colored by Seurat clustering, solid arrow indicating RHI/CTE depleted cluster. **c.** Stacked bar plot showing OPC Seurat cluster distribution across pathological groups. **d.** Bar plots showing OPC cluster distribution across control and pathological group or control and RHI-exposed samples, bar represents mean, error bars show standard error of the mean. Statistical analysis performed by ANOVA with Bonferroni correction (left) and two-tailed Mann-Whitney U test. **e.** Heatmap showing GO analysis of OPC cluster DEGs. **f.** UMAP showing oligodendrocytes colored by Seurat cluster, solid arrow indicates RHI and CTE depleted cluster. **g.** Stacked bar plot showing oligodendrocyte pathological group distribution per Seurat cluster. **h.** Bar plots representing cluster distribution across pathological groups or control and RHI-exposed samples. Bar represents mean, error bar represents standard error of the mean. Statistical analysis performed by ANOVA with Bonferroni correction (left) or two-tailed t-test (right). **i.** Heatmap showing GO analysis of oligodendrocyte cluster DEGs. **j.** UMAP showing T cells colored by Seurat cluster. **k.** Heatmap of GO analysis of T cell cluster DEGs.

**Supplementary Figure 2. Celda module workflow and cell type expression. a.** Celda module workflow diagram. **b.** Examples of Celda module expression in microglia. UMAPs show module expression, heatmaps show per-cell expression with genes listed on the right. Genes can also be viewed in Supplementary Table 17. **c.** Examples of Celda module expression in endothelial cells. UMAPs show module expression, heatmaps show per-cell expression with genes listed on the right. Genes can also be viewed in Supplementary Table 18.

**Supplementary Figure 3. Microglia Celda Modules.** Violin plots depicting Celda module expression for modules 1-90 across Homeostatic, RHIM1, RHIM2, and RHIM3 microglial clusters. Black bar is the median statistic from ggsignif.

**Supplementary Figure 4. Cellular subtype metadata correlations and Microglia external dataset projection. a.** Correlation heatmap of selected metadata and cellular subtypes. Heatmap color depicts and direction and magnitude of Pearson’s r correlation value. Statistical analysis performed by Pearson correlation. *, p <0.05, **, p < 0.01, ***, p < 0.001. **b.** UMAP of combined and reclustered microglia from Sun et al 2023 dataset and current dataset. Left, colored by microglial subtypes from Sun data, left colored by subtypes in current dataset.

**Supplementary Figure 5. Astrocytic responses to head trauma. a.** UMAP representing 4 astrocytic subtypes. **b.** UMAP from (a) colored by pathological group. **c.** Stacked bar plots showing astrocyte subtype distribution across pathological groups, statistics performed by chi-squared test. **d.** Bar plots showing astrocyte subcluster distribution in control and RHI-exposed samples, dots represent individual donors colored by pathological group identity. Bars represent mean, error bars represent standard error of the mean. Statistical analysis was performed using two-tailed Mann Whitney U-test. **e.** Stacked bar plots showing pathological distribution across astrocyte subtypes. **f.** Violin plots showing Celda module expression across astrocyte subtypes. Black bar showing median statistic. Colored by astrocyte subtype most associated with specific module expression. Statistical analysis performed by linear mixed effects model. **g.** Gene ontology analysis of astrocytic subtypes performed by Metascape. **h.** Dot plot representing expression of selected DEGs across astrocytic subtype and annotated by function.

**Supplementary Figure 6. Astrocyte Celda Modules.** Violin plots depicting Celda module expression for modules 1-80 across Astro1, Astro2, Astro3, and Astro4 astrocyte clusters. Black bar is the median statistic from ggsignif.

**Supplementary Figure 7. Endothelial Cell Celda Modules.** Violin plots depicting Celda module expression for modules 1-60 across Cap1, Cap2, Cap3, and Cap4 endothelial cell clusters. Black bar is the median statistic from ggsignif.

**Supplementary Figure 8. Neuronal layer subtype identification. a.** UMAP depicting all neurons clustered together colored by Seurat cluster. **b.** Dot plot of gene expression of inhibitory and excitatory neuron and astrocyte marker genes Seurat clusters from (a). **c.** UMAP from (a) colored by cell type determination. **d.** Stacked bar plot of sequencing batch distribution of Seurat clusters from (a). **e.** UMAP showing excitatory neurons colored by Seurat cluster. **f.** UMAP showing excitatory neurons colored by later subtype. **g.** Dot plot showing expression of excitatory neuron layer subtype genes in excitatory neuron Seurat clusters from (e). **h.** UMAP showing inhibitory neurons colored by Seurat cluster. **i.** UMAP showing inhibitory neurons colored by layer subtype. **j.** Dot plot showing expression of inhibitory neuron layer subtype genes across inhibitory neuron Seurat clusters from (h).

**Supplementary Figure 9. Neuron layer GO analysis, pathological group enrichment and RNAScope validation. a.** UMAP depicting excitatory neurons colored by layer subtype. **b.** Heatmap showing GO analysis of excitatory layer up and downregulated DEGs. **c.** Heatmap showing GO analysis of inhibitory layer up and downregulated DEGs. **d.** Bar plots of excitatory neuron layer proportions by pathological group. Bar represents mean, dots represent individual samples, error bars show standard error of the mean. Statistical analysis performed by ANOVA with Bonferroni correction. *, p<0.05, **, p<0.01. **e.** UMAP showing inhibitory neurons colored by layer subtype. **f.** Bar plots of inhibitory neuron layer proportions by pathological group. Bar represents mean, dots represent individual samples, error bars show standard error of the mean. Statistical analysis performed by ANOVA with Bonferroni correction. **g.** Representative image showing RNAScope *in situ* hybridization of CUX2/LAMP5 image analysis with correct anatomical layer-wise distribution. **g.** Representative image of CUX2/LAMP5 *in situ* with white squares showing HALO identification of double-positive cells.

## Supplementary Table Legends

**Supplementary Table 1. Demographics Summary of SnRNAseq Samples.** Table showing a summary of the demographic information for the snRNAseq samples. Data expressed as mean ± standard deviation. Age at death and years of exposure analyzed with one-way ANOVA.

**Supplementary Table 2. Demographics table of SnRNAseq Samples.** Table showing demographic information from samples included in snRNAseq dataset. PMI = post-mortem interval.

**Supplementary Table 3. Demographics table of *in situ* hybridization and Nissl Samples. a**. Table showing demographic information from samples included in *in situ* hybridization experiments. **b.** Table showing demographic information from samples included in Nissl staining experiments. PMI = post-mortem interval.

**Supplementary Table 4. List of Antibodies.**

**Supplementary Table 5. List of *In Situ* probes.**

**Supplementary Table 6. DEGs of All Cells in snRNAseq Dataset. a, b.** List of differentially expressed genes in respective clusters depicted in Figure 1c and Extended Data Figure 1l. Log2_FC = log-2 fold change. FDR = false discovery rate.

**Supplementary Table 7. DEGs of Excitatory and Inhibitory Neurons in snRNAseq Dataset. a.** List of differentially expressed genes in Seurat clusters of inhibitory and excitatory neuron clusters depicted in Supplementary Figure 2a. Log2_FC = log-2 fold change. FDR = false discovery rate.

**Supplementary Table 8. DEGs of Vascular Cells in snRNAseq Dataset. a.** List of differentially expressed genes in clusters depicted in Extended Data Figure 4a. Log2_FC = log-2 fold change. FDR = false discovery rate.

**Supplementary Table 9. DEGs of Microglia in snRNAseq Dataset. a.** List of differentially expressed genes in microglial subtype clusters depicted in Figure 2a. **b-e.** List of differentially expressed genes in microglial pathological group comparisons. **f.** List of genes significantly differentially expressed across pseudotime depicted in Figure 2l. Log2_FC = log-2 fold change. FDR = false discovery rate.

**Supplementary Table 10. DEGs of Astrocytes in snRNAseq Dataset. a.** List of differentially expressed genes in astrocyte subtype clusters depicted in Figure 4a. **b-e.** List of differentially expressed genes in endothelial pathological group comparisons. Log2_FC = log-2 fold change. FDR = false discovery rate.

**Supplementary Table 11. DEGs of Endothelial Cells in snRNAseq Dataset. a.** List of differentially expressed genes in endothelial subtype clusters depicted in Figure 3a. **b-e.** List of differentially expressed genes in astrocyte pathological group comparisons. Log2_FC = log-2 fold change. FDR = false discovery rate.

**Supplementary Table 12. DEGs of Oligodendrocytes in snRNAseq Dataset. a.** List of differentially expressed genes in oligodendrocyte Seurat clusters depicted in Supplementary Figure 1b. **b-e.** List of differentially expressed genes in oligodendrocyte pathological group comparisons. Log2_FC = log-2 fold change. FDR = false discovery rate.

**Supplementary Table 13. DEGs of Oligodendrocyte Precursor Cells in snRNAseq Dataset. a.** List of differentially expressed genes in oligodendrocyte Seurat clusters depicted in Supplementary Figure 1f. **b-e.** List of differentially expressed genes in oligodendrocyte precursor cell pathological group comparisons. Log2_FC = log-2 fold change. FDR = false discovery rate.

**Supplementary Table 14. DEGs of T Cells in snRNAseq Dataset. a.** List of differentially expressed genes in T Cell Seurat clusters depicted in Supplementary Figure 1j. Log2_FC = log-2 fold change. FDR = false discovery rate.

**Supplementary Table 15. DEGs of Excitatory Neurons in snRNAseq Dataset. a.** List of differentially expressed genes in excitatory neuron subtype clusters depicted in Figure 5a. **b-d.** List of differentially expressed genes in excitatory neuron pathological group comparisons **e-n.** List of differentially expressed genes in individual excitatory neuron subtype pathological group comparisons. Log2_FC = log-2 fold change. FDR = false discovery rate.

**Supplementary Table 16. DEGs of Inhibitory Neurons in snRNAseq Dataset. a.** List of differentially expressed genes in inhibitory neuron subtype clusters depicted in Supplementary Figure 2i. **b-d.** List of differentially expressed genes in inhibitory neuron pathological group comparisons **e-l.** List of differentially expressed genes in individual inhibitory neuron subtype pathological group comparisons. Log2_FC = log-2 fold change. FDR = false discovery rate.

**Supplementary Table 17. Microglia Celda Modules and LME statistics. a.** List of genes identified in each Celda module. **b.** Linear mixed effects model statistical output.

**Supplementary Table 18. Endothelial Cell Celda Modules and LME statistics. a.** List of genes identified in each Celda module. **b.** Linear mixed effects model statistical output.

**Supplementary Table 19. Astrocyte Celda Modules and LME statistics. a.** List of genes identified in each Celda module. **b.** Linear mixed effects model statistical output.

## Notes

### Competing Interest Statement

The authors have declared no competing interest.

### Summary of Updates

Added Celda gene module analyses to all cell types. Included in situ hybridization and protein staining for all cell types to validate genomic findings. Also, we included a new figure 5 which examines receptor-ligand pair interactions. We revised the majority of results to reflect this new data and findings.

