## Supplementary figures and images for "Repetitive head impacts induce neuronal loss and neuroinflammation in young athletes"

### Supplementary Figure 1

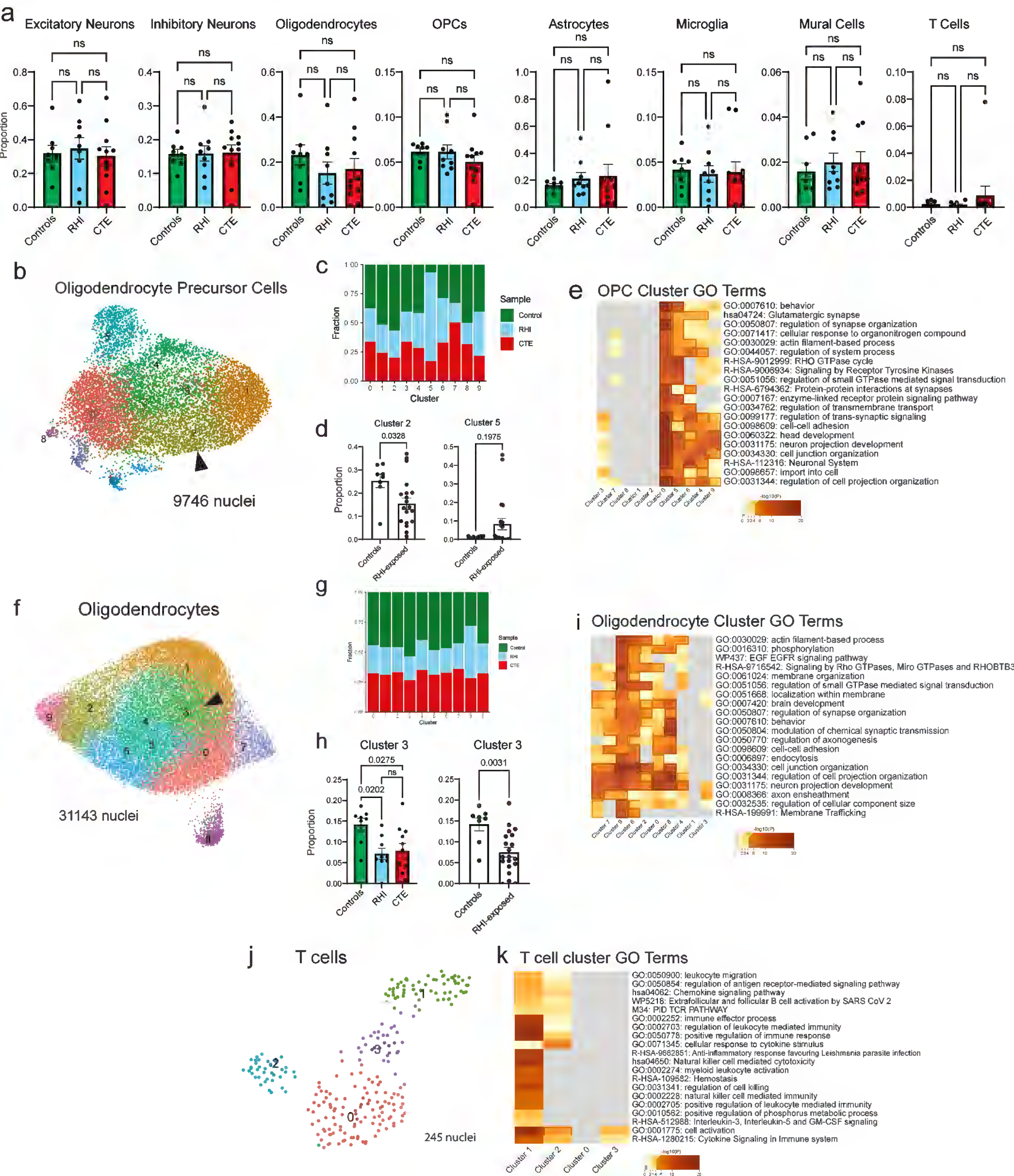

### Supplementary Figure 2

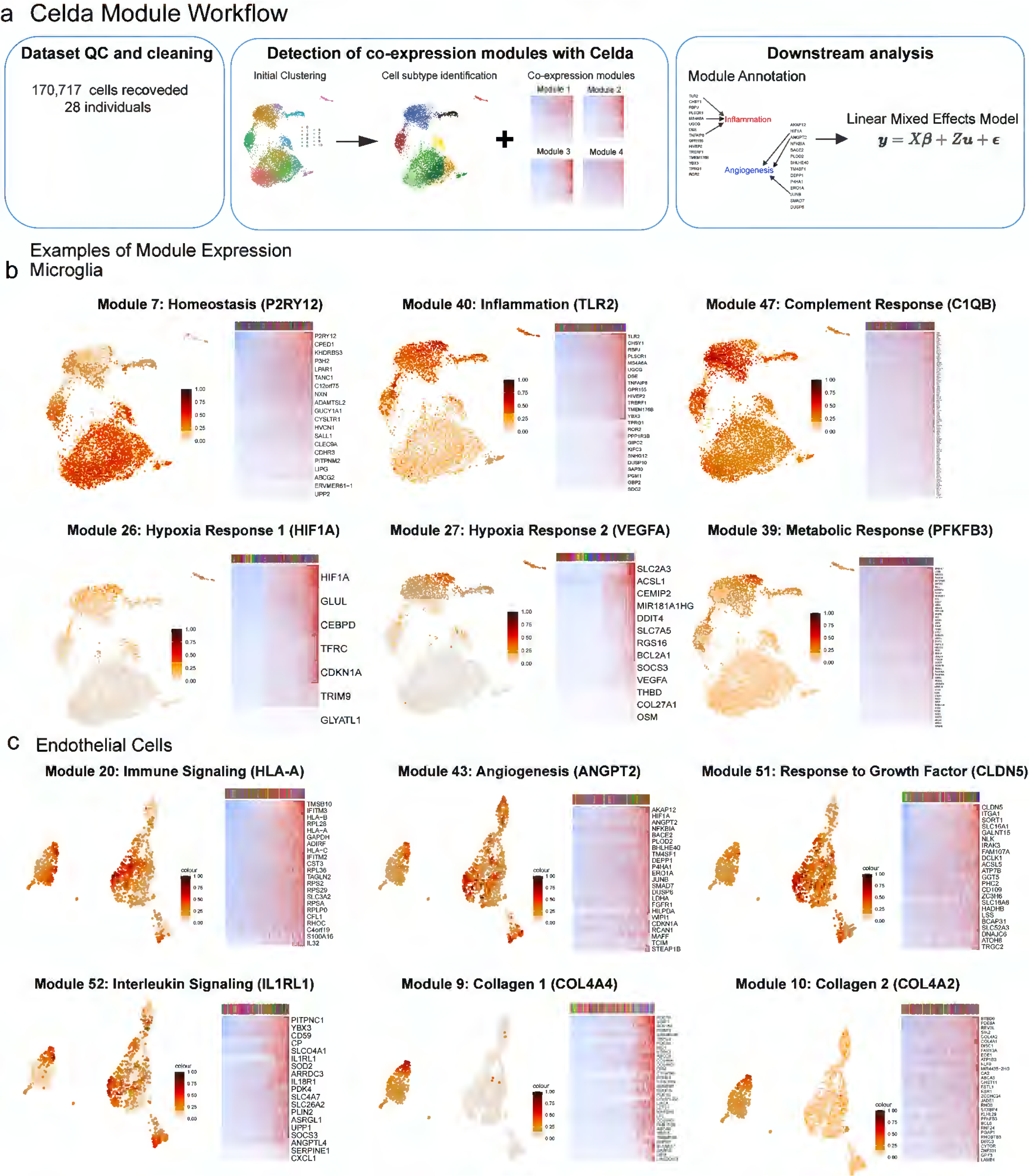

### Supplementary Figure 3

#### Microglia Modules L1- L90

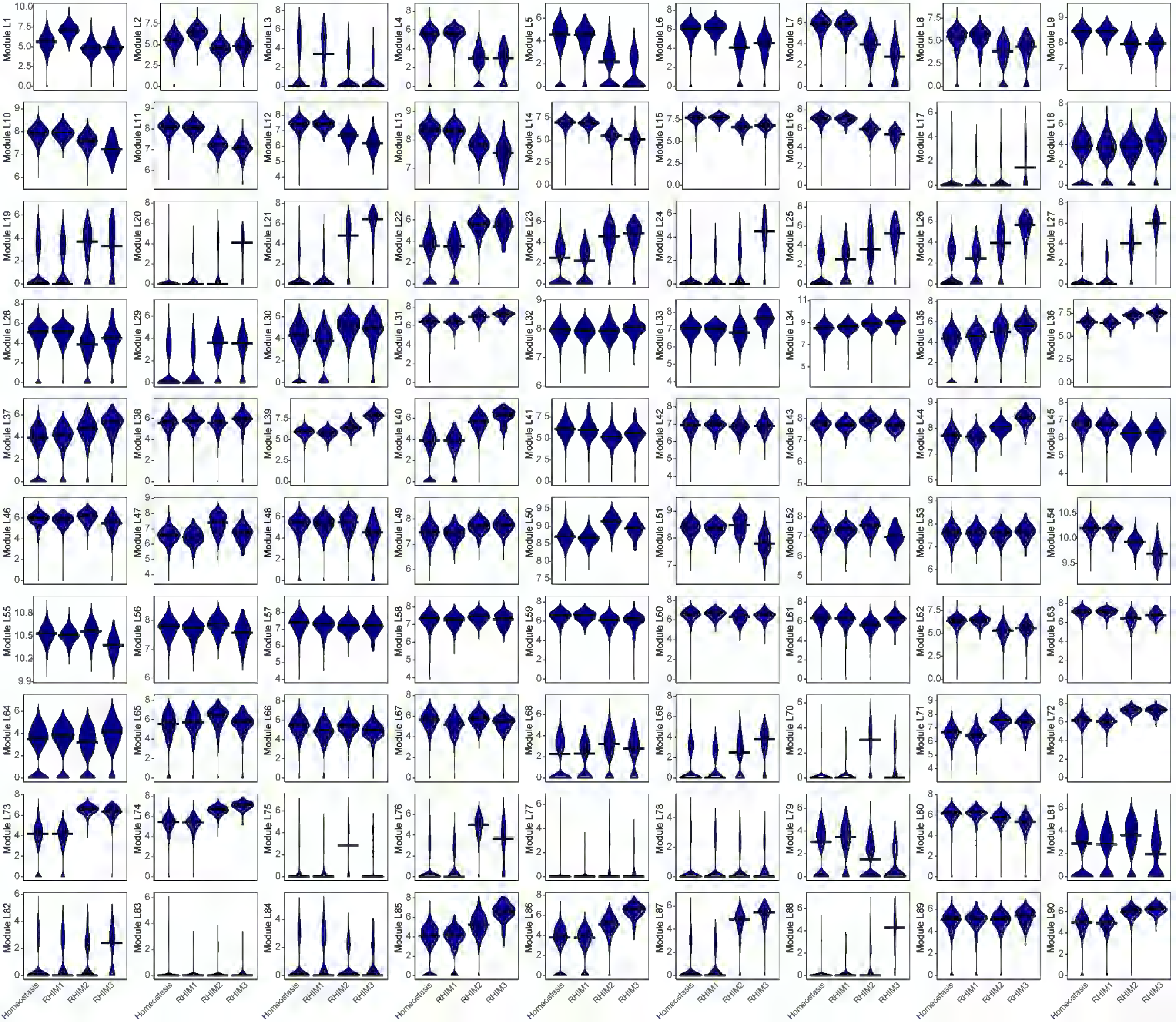

### Supplementary Figure 4

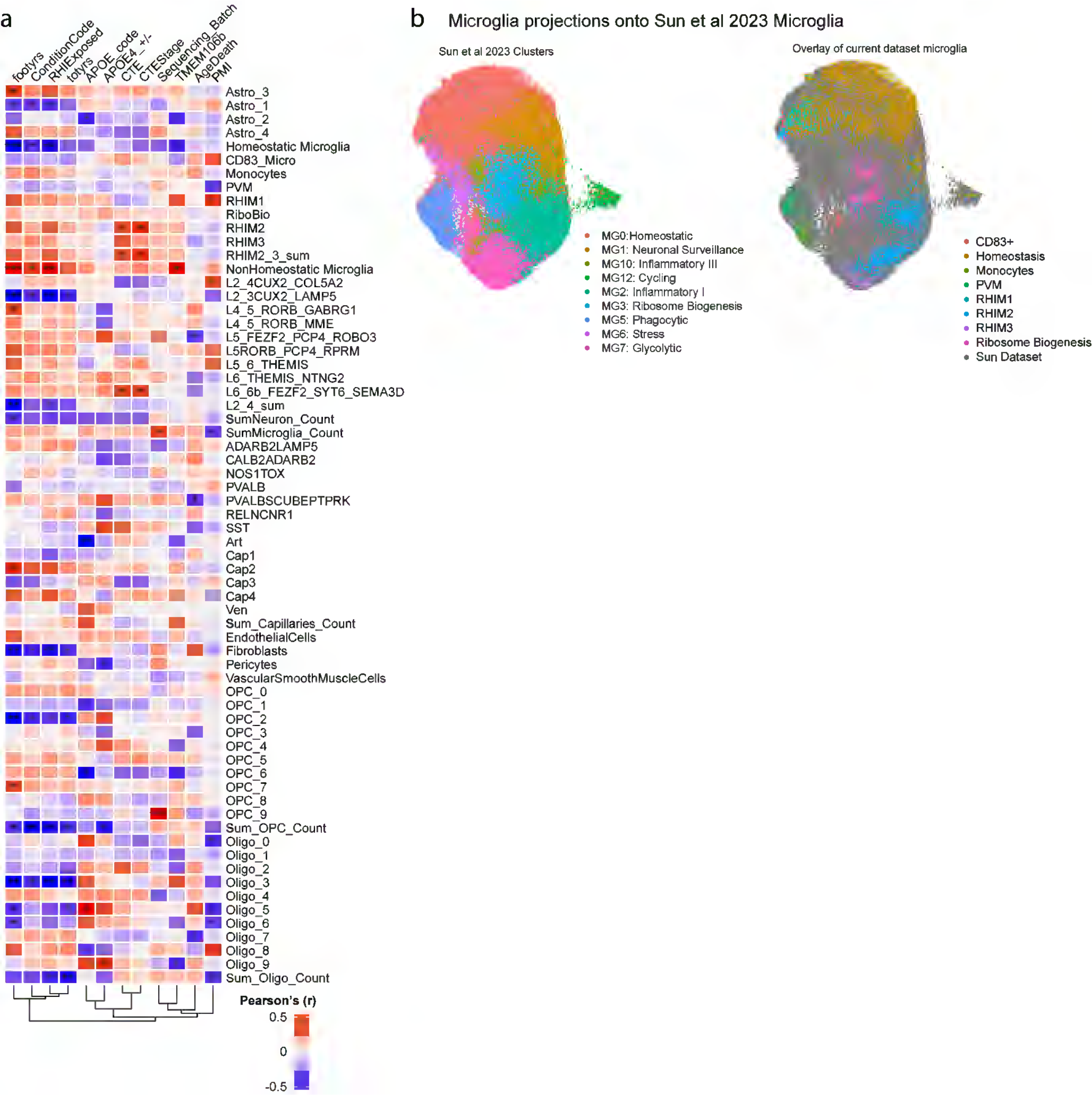

### Supplementary Figure 5

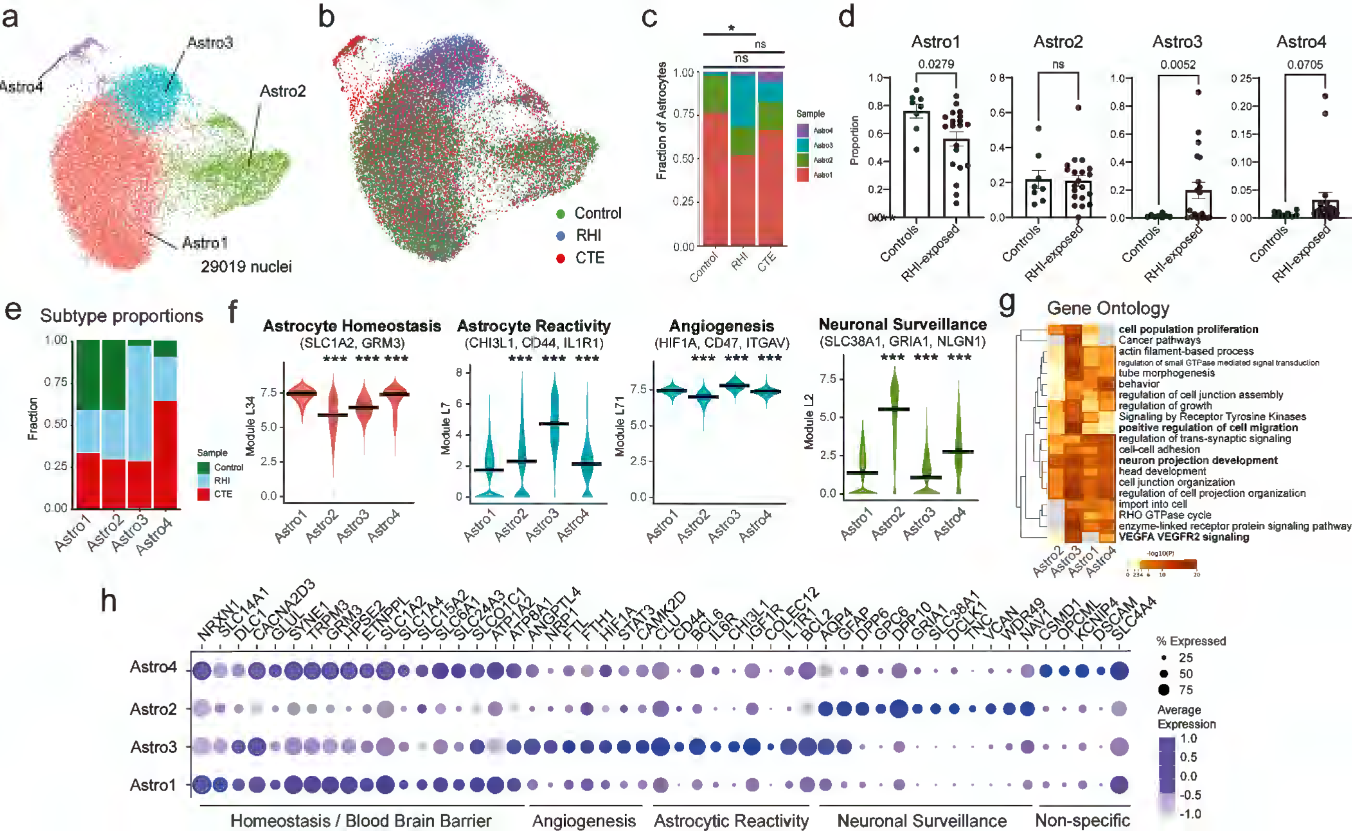

### Supplementary Figure 6

#### Astrocyte Modules L1-L80

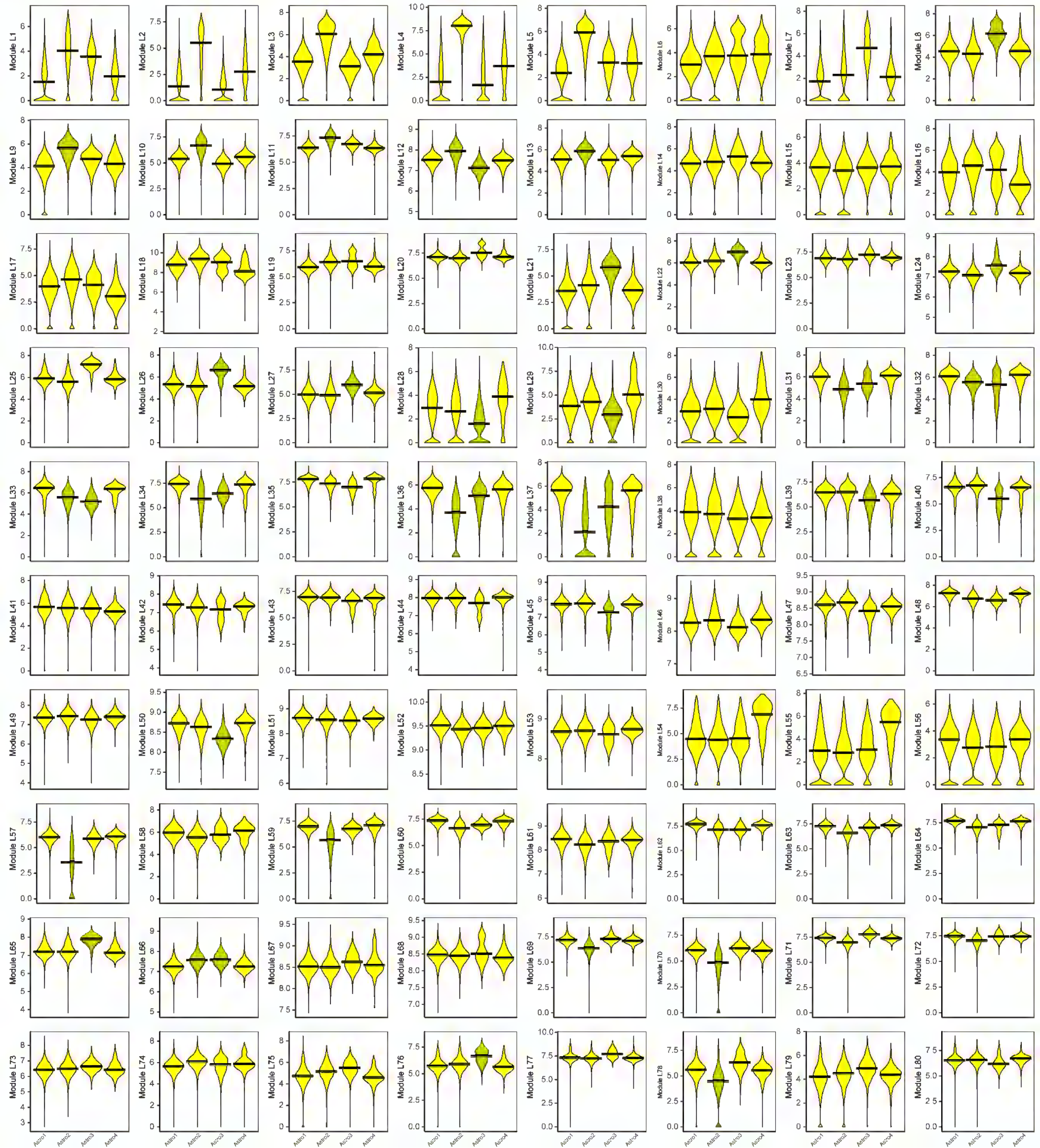

### Supplementary Figure 7

Endothelial Cell Modules L1- L60

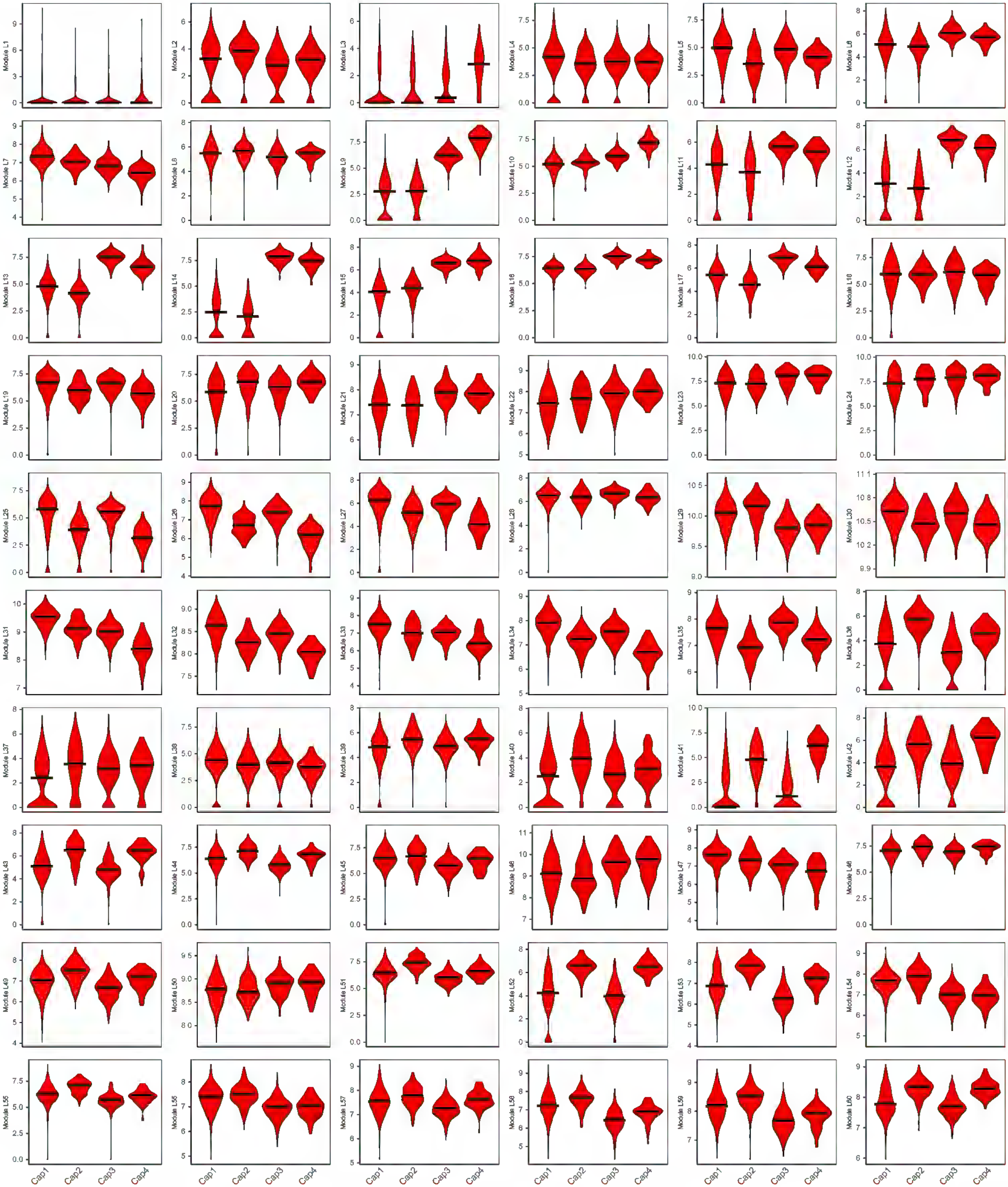

### Supplementary Figure 8

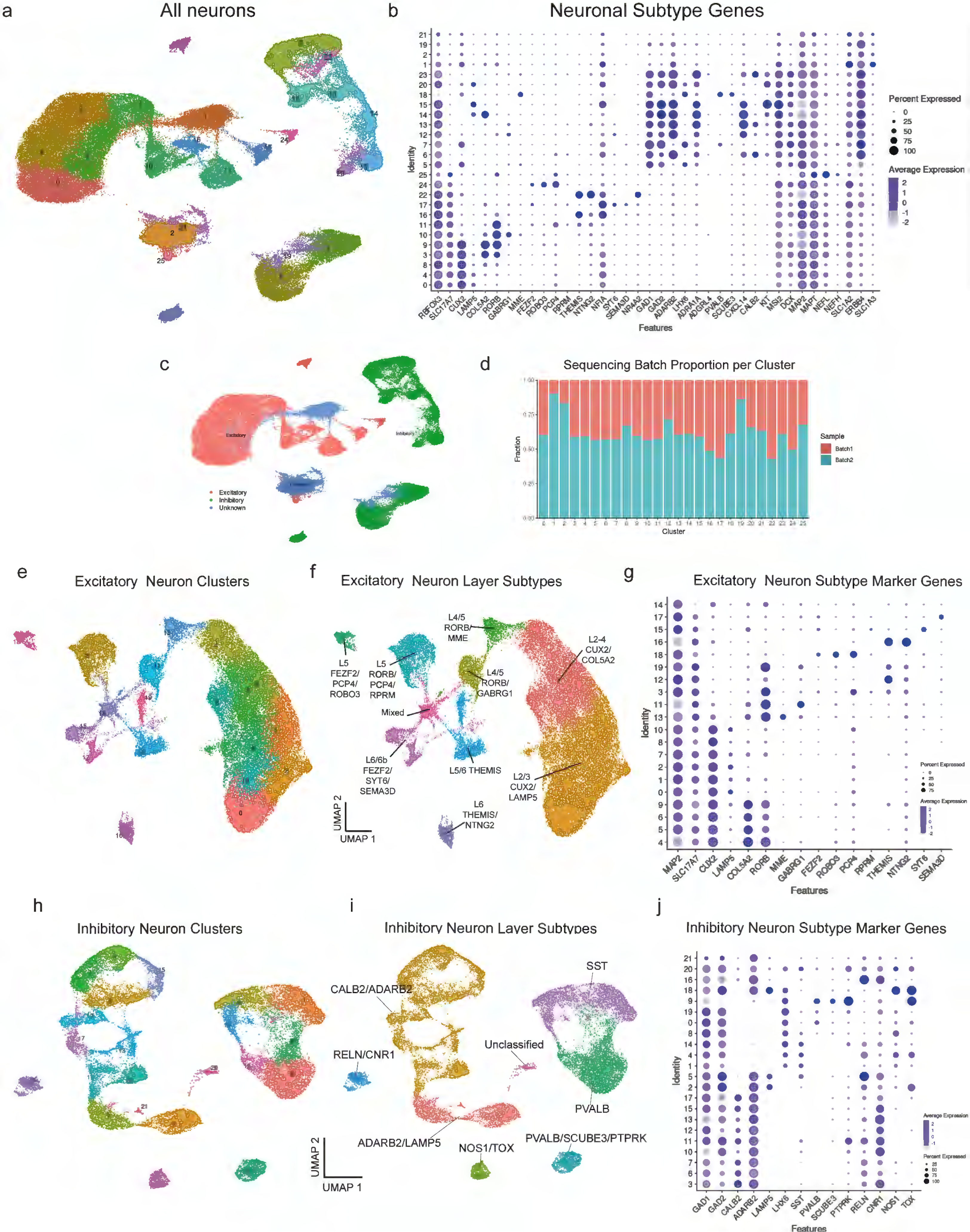

### Supplementary Figure 9

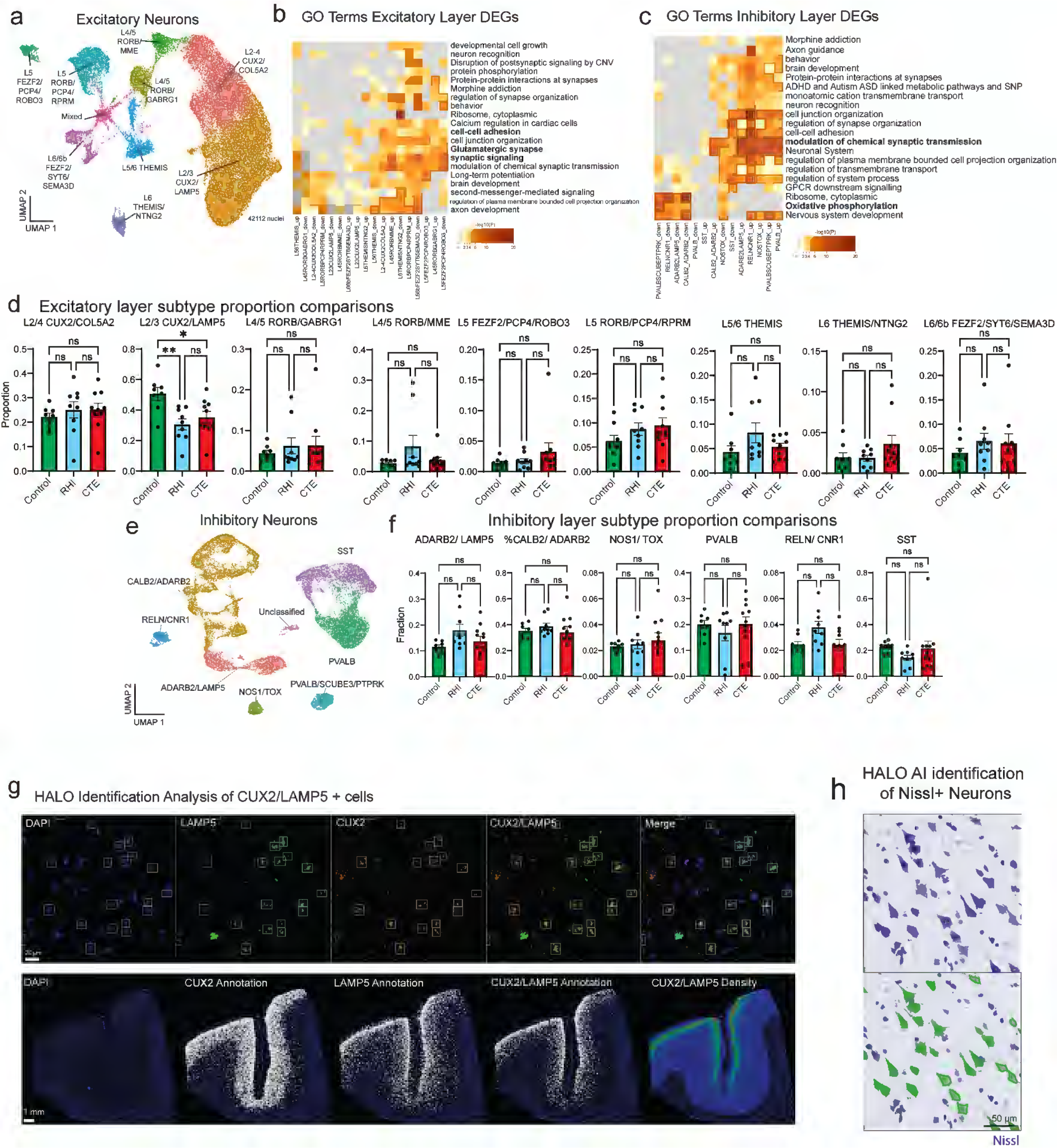
